# Spatial organization of enzymes and cofactors in synthetic membrane-associated condensates (sMACs)

**DOI:** 10.64898/2026.04.24.720695

**Authors:** Ka-Hei Siu, Virginia Jiang, Eoin M. Gaffney, Mackenzie T. Walls, Jerelle Joseph, José L. Avalos

## Abstract

Cells spatially organize metabolic enzymes to optimize flux through complex biochemical networks. Here, we engineered <u>s</u>ynthetic <u>M</u>embrane-<u>A</u>ssociated <u>C</u>ondensates (sMACs), artificial assemblies that form on intracellular membranes through liquid-liquid phase separation, to couple metabolic compartmentalization with cofactor partitioning. Using intrinsically disordered regions (IDRs) characterized by simulations and validated by experiments, we identified an IDR sequence, Dbp1n, that selectively enrich NADH and NADPH, establishing distinct local redox microenvironments. At the endoplasmic reticulum (ER) membrane, NADPH-enriched sMACs enhanced cytochrome P450 activity by improving electron transfer and supporting *in situ* regeneration. At the plasma membrane, NADH-enriched sMACs provided scaffolds for a xylose uptake and assimilation pathway that exploits localized NADH/NAD⁺ recycling to improve flux. These results collectively demonstrated that condensate composition and membrane context can be engineered to enhance metabolic functions. sMACs thus provide a modular framework for creating programmable, membrane-associated microreactors, enabling new strategies for metabolic engineering and offering insights into how phase separation can be exploited in living cells.

## Main

Living cells must organize metabolic reactions in both space and time to ensure efficient flux through complex biochemical networks. One evolutionary strategy to meet this requirement is compartmentalization, which occurs either within membrane-bound organelles, such as mitochondria or the endoplasmic reticulum (ER), or in membrane-less biomolecular condensates formed through liquid–liquid phase separation (LLPS)^1^. Biomolecular condensates have emerged as versatile regulators of metabolism, capable of accelerating flux by concentrating enzymes and intermediates into dynamic assemblies known as metabolons^2^. A well-known example is the glycolytic body (G-body), in which enzymes of glucose metabolism form a condensate that enhances turnover to pyruvate and oxaloacetate^3–5^. These assemblies are not static but instead adapt to cellular states; for instance, purinosomes form to cluster enzymes for purine biosynthesis when cells have a high demand for purines^6^. Similarly, the G-body in *Saccharomyces cerevisiae* assemble glycolytic enzymes into a higher-order condensate, the “META body” (Metabolic Enzymes Transiently Assembling), under hypoxic conditions^7,8^. These examples highlight two defining features of condensates in metabolic regulation: 1) the ability to spatially organize enzymes to promote “substrate proximity channeling”^9^ at specific locations within the cell and 2) the capacity to temporally reconfigure metabolic flux in response to environmental or cellular demands.

Biomolecular condensates have further been shown to assemble on the surfaces of membrane-bound organelles, here referred to as “membrane-associated condensates.” Native membrane-associated condensates serve diverse roles in a cell, which include plugging damaged membranes^10^, regulating membrane morphology^11,12^, sequestering specific proteins^13,14^, and concentrating small molecules of interest^15^. One notable example is a glycolytic metabolon that localizes to the plasma membrane of pancreatic α/β cells in humans and mice^16^. In this system, the upper glycolytic pathway regulates the activity of an ATP-sensitive potassium ion channel, while the lower glycolytic pathway couples redox reactions that locally recycle NAD⁺/NADH to maintain cofactor balance. This example shows how membrane association and cofactor regulation work in tandem to promote metabolic flux by coupling spatial organization with local biochemical environments that sustain enzymatic activity.

In metabolic engineering applications, such coupling would be especially valuable because biosynthetic pathway flux depends on both substrate and cofactor availability. Colocalizing enzymes and transporters within condensates could increase rates of collision between reactants and catalysts locally while reducing losses to diffusion. Condensates can also facilitate substrate proximity channeling, enabling more efficient transfer of intermediates between successive enzymes in a cascade^9^. However, controlling metabolic flux alone is often insufficient, as many pathways of interest also require redox balance^17^. Existing approaches to maintain intracellular redox homeostasis include deleting competing pathways^18^, introducing enzymes that regenerate consumed cofactors^19^, engineering mutant enzymes with altered cofactor specificity^19,20^, or recycling cofactors consumed in one step of a pathway by regenerating them in subsequent steps^21,22^. These strategies highlight the importance of regulating both spatial organization and redox dynamics to maximize productivity.

Inspired by these principles, we hypothesized that engineered condensates on intracellular membranes could provide a modular means to control both substrate flux and redox balance for metabolic engineering applications. Accordingly, we sought to develop a new class of scaffolds that not only bring enzymes together but also actively shape their local spatial and biochemical environment, focusing on adherence to cellular membranes and co-factor recruitment. We adopted a synthetic biology approach to create artificial membrane-associated biomolecular condensates in *Saccharomyces cerevisiae*, which we call <u>s</u>ynthetic <u>M</u>embrane-<u>A</u>ssociated <u>C</u>ondensates (sMACs). These condensates can selectively sequester enzymes, substrates, and cofactors to amplify chemical potentials and drive reactions forward, overcoming transport bottlenecks frequently encountered on intracellular membrane surfaces. These sMACs assemble as film-like structures and can recruit cargo proteins, effectively functioning as synthetic compartments that spatially organize proteins to perform designed functions. We specifically demonstrated that metabolic reactions of biotechnological interest can be localized within sMACs. Furthermore, to explore how these condensates might modulate local redox balance, we developed a computational model to investigate how recycling cofactors could facilitate metabolic channeling between colocalized enzymes, thereby increasing flux to product by minimizing diffusive losses. We envision sMACs as versatile platforms not only for engineering new metabolic functions in diverse metabolic engineering applications but also for probing the fundamental principles that govern condensate formation and biochemical properties near cellular membranes.

## Results

### Design and construction of membrane-associated condensates

The basic building block of sMACs is a chimeric fusion protein consisting of at least two modular domains: 1) an intrinsically disordered region (“IDR”) containing multiple self-associating motifs that promotes weak, multivalent interactions to drive LLPS, such as those commonly found in nucleic-acid-binding proteins like Dbp1p from *S. cerevisiae*^23^ or FUS^24^, and 2) a membrane targeting region containing signals for trafficking the chimeric protein to the intended organelle and anchoring it to the membrane surface (Fig 1a). This basic construct may additionally contain one or more protein–protein interaction domains (PpIDs)^25^ to recruit cargo proteins to functionalize condensates (see below), as well as a fluorophore to visualize them with fluorescence microscopy (Supplementary Note 1).

**Figure 1.**
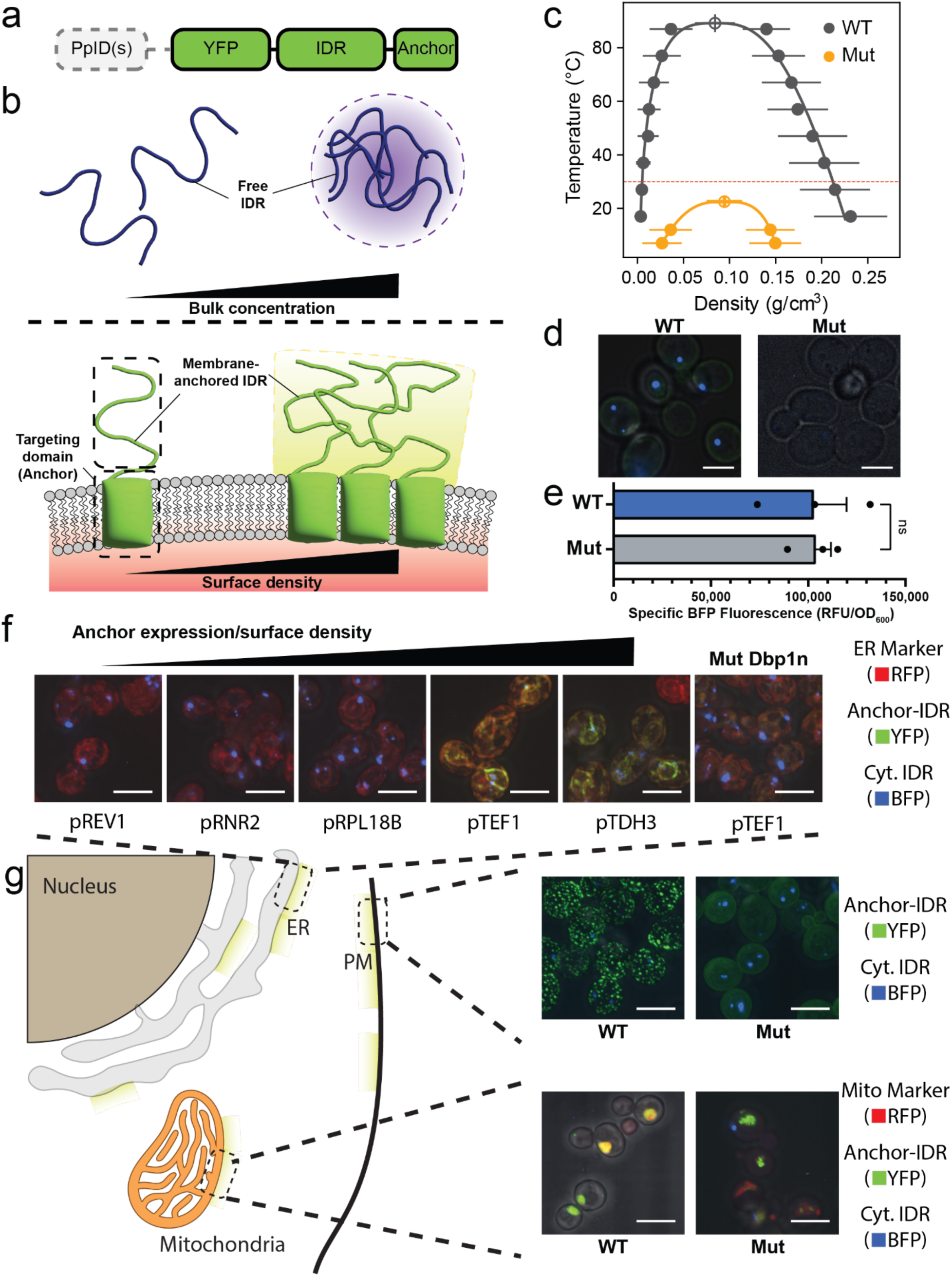
Design and construction of synthetic membrane-associated condensates (sMACs). **(a)** Schematic of sMAC basic building block, which consists of a membrane-targeting domain (“Anchor”), an intrinsically disordered region (“IDR”), and a Venus yellow fluorescent protein marker (“YFP”). One or more protein-protein interaction domains (“PpID(s)”) can be added to recruit specific cargo into sMACs. **(b)** Some IDRs with weak, multivalent interaction motifs can spontaneously demix from the surrounding solution and condense into a dense phase in the right conditions and at sufficiently high concentrations. Analogously, when an IDR is fused to a membrane-targeting domain and expressed at sufficiently high surface density, IDR-driven phase separation occurs on the targeted membrane surface, forming a synthetic condensate. **(c)** Computed phase diagrams (errors obtained through block-averaging) showed upper critical solution temperature (UCST) above optimal *S. cerevisiae* growth temperature (30°C, denoted by red dotted line) for Dbp1n (“WT”) but subcritical for Mut Dbp1n (“Mut”). Points and error bars represent the mean and standard error calculated from block-averaged simulation. **(d)** Expressing BFP-labeled cytosolic trimeric coiled coil-Dbp1n formed fluorescent puncta in yeast but not when Mut Dbp1n was used in place of WT Dbp1n despite **(e)** similar levels of expression as measured by BFP fluorescence, validating the predictions from the computed phase diagrams. **(f)** Expressing a YFP-labeled ER-targeting IDR fusion protein using sufficiently high strength constitutive promoters (pTEF1 and pTDH3) resulted in formation of fluorescent puncta (shown in green) on ER membrane (labeled with RFP and shown in red). The BFP-labeled cytosolic trimeric coiled-coil-Dbp1n fusion protein was co-expressed and only merged with the YFP-labeled ER puncta when ER-anchored Dbp1n was expressed at a sufficiently high level but remained cytosolic at either lower expression levels or when Mut Dbp1n was used, suggesting the YFP-labeled ER puncta were miscible liquid-like structures. **(g)** YFP-labeled puncta also formed on the plasma membrane or the outer membrane of the mitochondria (labeled with RFP and shown in red) when the corresponding membrane targeting domain was substituted into the base unit (Gap1p and Fis1p respectively). Note the lack of distinct green puncta structures when Mut Dbp1n was used in place of WT Dbp1n. All scale bars represent 5 µm. Two-tailed Student’s *t*-test was used to compare the sample data between each group of three biological replicates. All data are shown as the mean ± s.e.m. ns denotes not significant.

Like IDRs capable of condensing in bulk solution, the primary sequence of membrane-anchored IDR dictates both the critical parameters (e.g. concentration, temperature) at which phase separation occurs (Fig 1b)^24,26^ and the physicochemical properties (e.g. pH, partition coefficients)^1,27^ of the resulting membrane-associated condensate. To ensure that the membrane-anchored IDR retains the ability to condense in *S. cerevisiae* at its optimal growth temperature, we employed the Mpipi coarse-grained forcefield^28^ to run direct coexistence molecular dynamics (MD) simulations of the N-terminal region of Dbp1p (aa 2-154, “Dbp1n”, Supplementary Table 1). This IDR has recently been shown to modulate NADH partitioning *in vitro*^23^, which would be useful when compartmentalizing pathways that require this cofactor. We found that Dbp1n has an upper critical solution temperature (UCST) over 85°C (Fig 1c, “WT”, dark grey), well above the growth temperature of *S. cerevisiae*. Consistent with this result, expression of Dbp1n, tagged with a blue fluorescent protein (BFP) and a trimeric coiled-coil domain to promote condensate nucleation (Supplementary Note 2, Supplementary Fig 1), forms distinct blue puncta in baker’s yeast (Fig 1d, “WT”, left panel). To investigate the sequence features responsible for Dbp1n condensation, we applied a machine-learning driven bioinformatics pipeline to identify repeated motifs enriched in a library of intrinsically disordered proteins (See Methods and Extended Data Fig 1a) that likely undergo phase separation. According to the Mpipi-based IDR interaction predictor FINCHES^29^, key motifs identified within Dbp1n exhibit favorable energies of self-interaction, likely contributing to its propensity for phase separation (Extended Data Fig 1b).

To test this hypothesis, we designed a mutated variant of Dbp1n (“Mut Dbp1n”) that preserves the same fraction of charged residues, sequence patterning, and overall length as Dbp1n but is not expected to phase separate at similar conditions. Mut Dbp1n contains substitutions within the identified motifs, replacing residues known to be involved in interactions leading to LLPS for residues known to disrupt self-association required for phase separation^30–32^ (Supplementary Table 1). We performed the same direct coexistence MD simulations on Mut Dbp1n and found that the mutated variant fails to undergo phase separation at optimal yeast growth temperature, as its UCST lies lower than 30°C (Fig 1c, “Mut”, orange). These computational results were validated experimentally by expressing Mut Dbp1n, again tagged with BFP and a trimeric coiled-coil domain, in yeast cytosol. While the wild type Dbp1n forms distinct blue puncta (Fig 1d, left panel), Mut Dbp1n fails to make these condensates at the same expression levels and conditions (Fig 1d, “Mut”, right panel; Fig 1e).

Analogous to bulk concentrations required to achieve phase separation, we hypothesized that sMACs will only form at sufficiently high surface density. To test this on an intracellular membrane, we fused Dbp1n to the transmembrane domain of the endoplasmic reticulum (ER)-resident ubiquitin-conjugating enzyme Ubc6p (aa 233-250) and a Venus yellow fluorescent protein (YFP) marker. As an independent condensate marker, we expressed a trimeric coiled-coil–BFP–Dbp1n construct using the medium strength constitutive promoter pHHF2, which forms cytosolic condensates that merge onto the ER membrane only when sMACs assemble at the ER surface (Extended Data Fig 2a). Gradual increases in the expression level of ER-anchored Dbp1n, and hence its effective surface concentration, led to coalescence of wild-type Dbp1n–YFP and BFP puncta into discrete patches forming on the surface of the RFP-labeled ER (Fig 1f and Extended Data Fig 2b). This threshold was reached when YFP-Dbp1n was anchored to the ER and expressed using the medium-high strength constitutive promoter pTEF1. In contrast, expressing YFP-Mut Dbp1n using the same ER anchor and promoter produced only faint, diffuse YFP signals that did not colocalize with the cytosolic BFP–wild-type Dbp1n puncta (Fig 1f, far right panel). These results demonstrate that membrane-anchored Dbp1n drives phase separation at sufficiently high surface densities and that sMACs remain miscible with cytosolic non-anchored IDRs, a key indicator of liquid-like behaviors of biomolecular condensates as opposed to solid-like structures resulting from protein aggregation.

sMACs can be assembled on other intracellular membranes using the same pTEF1 promoter and expression cassette by replacing the ER-targeting Ubc6p(233-250) domain with membrane-specific anchors, such as the plasma membrane (PM)-targeting Gap1p (aa 550-602)^33^ or the mitochondrial outer membrane-targeting Fis1p (aa 119-155)^34^ domains (Fig 1g). This ability to redirect membrane-anchored IDRs to distinct organelles highlights the modularity of the design and suggests that the sMAC platform can be flexibly deployed across different subcellular contexts.

### Engineered recruitment of protein into membrane-associated condensates

To functionalize sMACs, we incorporated engineered protein–protein interaction domains (PpIDs) into the basic building block design (Fig 2a). These modular domains enable selective recruitment of protein cargo into condensates, with recruitment efficiency governed by the apparent affinity between each domain and its cognate ligand^25^. Because these PpIDs are orthogonal^25^, different combinations can be assembled onto the same scaffold to achieve simultaneous recruitment of multiple cargo proteins with independently tunable stoichiometries^35^. This modular design provides a mechanism to control local enzyme ratios within sMACs, thereby facilitating fine-tuning of pathway flux within the synthetic compartments.

**Figure 2.**
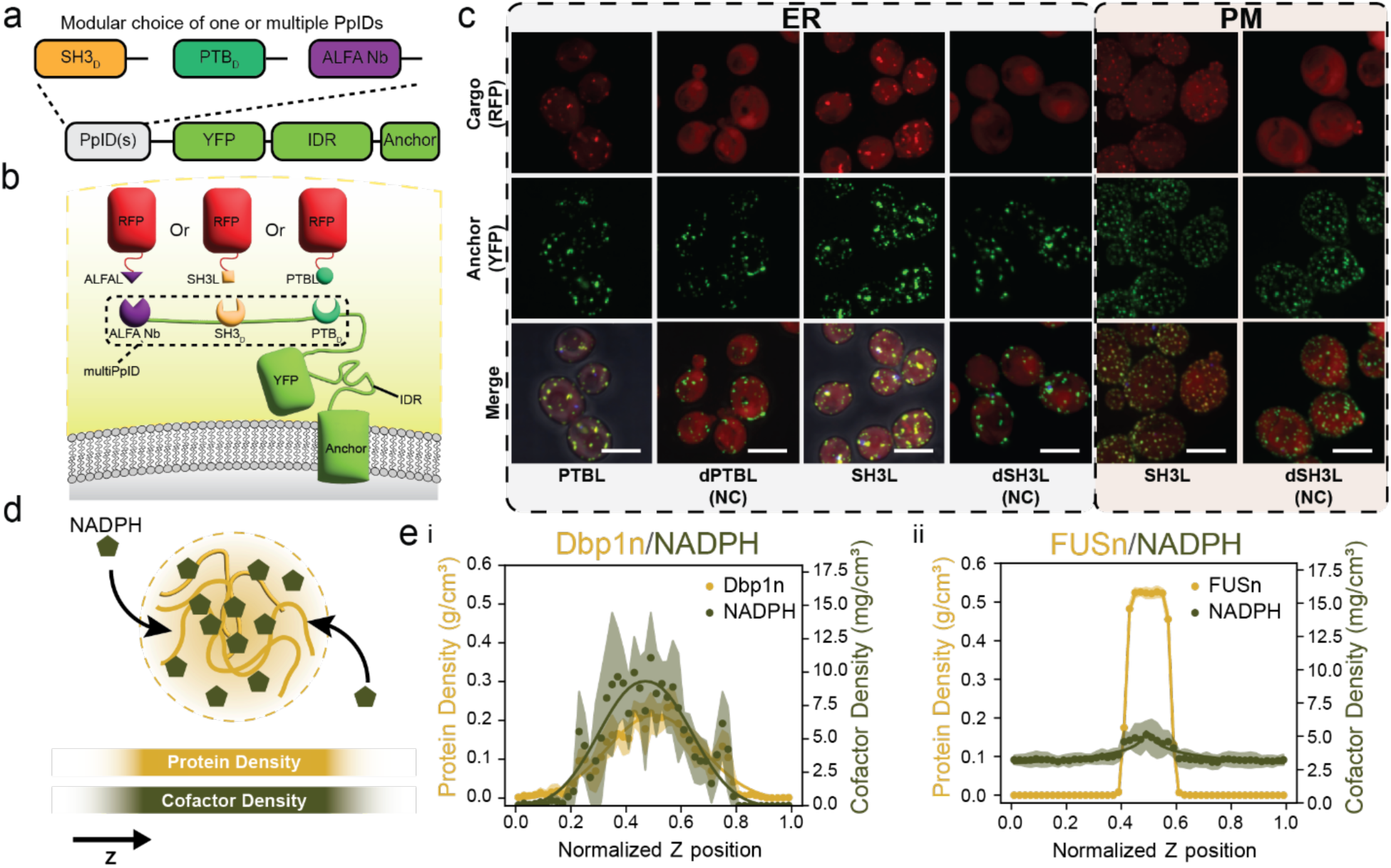
Protein-protein interaction domain (PpID)-dependent recruitment of cargo proteins and partition of cofactors to functionalize sMACs. **(a)** One or more PpIDs can be fused to the basic building block of sMACs to enable **(b)** recruitment of cargo proteins fused to the cognate peptide ligands (e.g. PTBL, SH3L, or ALFAL). Use of multiple PpIDs (“multiPpID”) allows simultaneous orthogonal recruitment of specific cargo proteins into sMACs. **(c)** Representative microscopy images of C-terminally tagged RFP-peptide ligand cargo recruitment to ER-associated sMACs with each column representing a different peptide ligand denoted at the bottom of each column. Merged images show cargo in red, and multiPpID-YFP-Dbp1n-Anchor in green. “NC” denotes negative control strains where the corresponding ligand (“dPTBL” and “dSH3L”) is inactivated and incapable of binding by point mutations in the peptide sequence. All scale bar represent 5 µm. **(d)** Small molecules such as the reducing cofactor NADPH may selectively partition into condensates, depending on interactions between the IDR and the molecules. **(e)** i: NADPH partitioned into the dense phase of Dbp1n condensates in coarse-grained direct coexistence molecular dynamics simulation. ii: In FUSn condensates, NADPH was weakly enriched in the dense phase over the dilute phase in coarse-grained direct coexistence molecular dynamics simulation. Points represent the time-averaged density over equilibrated simulation frames at a given Z-axis position, shaded region represents the standard error, and curve represents a super-Gaussian fit to data.

Accordingly, we created “multiPpID” sMAC building blocks that incorporate ALFA nanobody, Crk SH3, and PTB domains into a single construct, and tested whether each domain retained the ability to recruit its intended cargo (Fig. 2b, Supplementary Note 1). Using mRuby2 RFP fused to the cognate peptide ligand for the three PpIDs, which span a range of binding affinities (Extended Data Table 1), we demonstrated efficient and specific cargo recruitment into sMACs targeted to the ER and plasma membrane (Fig 2c and Extended Data Fig 3). Interestingly, the fusion of PpIDs to the sMAC scaffold slightly alters condensate morphology, possibly due to the inherent changes in its overall composition (Fig 2c and Extended Data Fig 3). Nonetheless, all tested PpIDs successfully mediated selective cargo recruitment, indicating that protein loading into sMACs can be achieved and predictably tuned by the PpID affinity as previously shown^25^. This modular control enables the design of membrane-localized compartments capable of concentrating selected enzymes or complexes for targeted metabolic processes.

### Co-factor partitioning into condensates depends on IDR properties

Beyond recruiting proteins to condensates using PpIDs, we sought to promote the selective partitioning of small molecules, specifically metabolic cofactors such as NADPH, into sMACs (Fig 2c). The ability to design the biochemical environment in synthetic condensates would be an empowering strategy to support heterologous metabolic pathways. *In vitro* studies have shown that small molecules can partition differentially between the dense and dilute phases of condensates, a behavior largely governed by interactions between the small molecules and the IDRs that drive condensate formation^23,27,36,37^.

Previous work has shown that sets of homologous IDRs can be identified using protein alignment-based methods applied to adjacent folded domains^38,39^. We sought to uncover the molecular basis for previously described cofactor partitioning in Dbp1n condensates by performing a bioinformatic search for ATP-dependent RNA helicases similar in function to Dbp1p. From this sequence database, we identified conserved motifs across the library of IDRs. We reasoned that some of these motifs may be implicated in NADH partitioning through formation of favorable enthalpic interactions with NADH. As we had hypothesized, yeast Dbp1n contains many of the motifs (Extended Data Table 2) common to ATP-dependent helicases and has key properties implicated in phase separation propensity and binding of charged nucleotides such as low radius of gyration and high fraction of charged residues (Extended Data Figure 4a). By contrast, the N-terminal sequence of the human RNA-binding protein FUS (aa 1-214, “FUSn”, Supplementary Table 1) lacks the most frequently identified sequence motifs in ATP-dependent helicases (Extended Data Table 2).

Having established sequence-level differences between Dbp1n and FUSn, we next investigated whether these differences translate into differential cofactor partitioning. To examine this *in silico*, we parameterized NADH, NAD⁺, NADPH, and NADP⁺ for residue-resolution coarse-grained molecular dynamics. To investigate affinity of the Dbp1n IDR motifs for NADH, we ran single chain Dbp1n simulations in the presence of a molar excess of NADH and found that the identified motifs, especially Motif 5, were present in the regions of the protein with highest frequency of forming NADH contacts (Extended Data 4b-c). In order to extend these observations to collective phase behavior, we then performed direct coexistence simulations at 30 °C with Dbp1n, Mut Dbp1n, and FUSn. All four cofactors partitioned more strongly into Dbp1n condensates compared to FUSn condensates (Fig 2d, Extended Data Fig 5, Table 1, and Supplementary Video 1 and 2). To experimentally validate NAD(P)H partitioning predictions, we purified Dbp1n fused to mCherry and a trimeric coiled-coil domain (see Methods) and measured the NAD(P)H fluorescence of the dense and dilute phases after mechanical separated by centrifugation (see Methods, Extended Data Figure 6a). The experimentally measured partition coefficients were in close agreement with those obtained from simulation (Extended Data Fig 6b,c and Table 1). The agreement between simulation and experimental results suggests that our computational framework can be extended to validate and predict the partitioning behavior of additional cofactors in both native and engineered biomolecular condensate systems, including sMACs.

**Table 1.** Partition coefficients (K_p_) for NADPH/NADP^+^ and NADH/NAD^+^ in condensates formed with different IDRs estimated from computational or experimental methods.

| IDR | Cofactor | Type* | $K_p^{\wedge}$ | ± s.e.m.<sup>#</sup> |
| --- | --- | --- | --- | --- |
| Dbp1n | NADPH | Slab | 27.50 | ± 3.74 |
| | | Exp | 35.54 | ± 3.84 |
| | | Mem | 13.07 | ± 5.08 |
| | NADP<sup>+</sup> | Slab | 11.55 | ± 2.87 |
| | | Mem | 7.66 | ± 1.92 |
| | NADH | Slab | 8.82 | ± 2.32 |
| | | Exp | 8.65 | ± 1.93 |
| | | Mem | 19.53 | ± 5.91 |
| | NAD<sup>+</sup> | Slab | 7.17 | ± 2.00 |
| | | Mem | 3.29 | ± 0.68 |
| FUSn | NADPH | Slab | 1.44 | ± 0.05 |
| | | Mem | 1.45 | ± 0.34 |
| | NADP<sup>+</sup> | Slab | 1.52 | ± 0.64 |
| | | Mem | 2.68 | ± 1.05 |
| | NADH | Slab | 2.37 | ± 0.87 |
| | | Mem | 6.03 | ± 1.49 |
| | NAD<sup>+</sup> | Slab | 1.90 | ± 0.60 |
| | | Mem | 2.23 | ± 0.80 |
| Mut Dbp1n | NADPH | Slab | 0.91 | ± 0.06 |
| | NADP<sup>+</sup> | Slab | 1.28 | ± 0.06 |
| | NADH | Slab | 1.02 | ± 0.05 |
| | NAD<sup>+</sup> | Slab | 4.05 | ± 1.21 |

\*Type denotes estimate from either 1) “Slab”: MD simulations performed in slab system, or 2) “Exp”: experimental results measured via fluorescence comparing the dense and dilute phases, or 3) “Mem”: MD simulations performed in membrane-tethered system.
<sup>^</sup> $K_p$ denotes the sample mean of the partition coefficient
<sup>#</sup>s.e.m. denotes standard error of the mean

Taken together, these results show that sMACs can be not only functionalized to enhance local enzyme concentrations but also designed to increase local concentrations of small molecules such as cofactors and potentially substrates or intermediates by using appropriate IDR sequences. Accordingly, sMACs can be used to create local biochemical environments to drive metabolic reactions forward, potentially beyond what is achievable in the cytosolic milieu due to enhancement of the local cofactor concentration. These potential advantages are showcased by the examples described below.

### ER-associated synthetic metabolon designed to enhance cytochrome P450 productivity

An important class of enzymes that can benefit from sMACs localization is the eukaryotic cytochrome P450 enzymes (CYPs), which localize mainly to the endoplasmic reticulum (ER)^40^. These membrane-bound enzymes play key roles in various metabolic processes, including drug metabolism in humans^41^ and the biosynthesis of plant natural products^42^. CYPs often represent a major bottleneck in natural product biosynthesis for microbial cell factories^43^; yet, they are indispensable for introducing oxygenated functional groups that enhance structural diversity and biological function^42^. Their efficiency is frequently limited by poor expression and low catalytic activity caused by inefficient electron transfer between CYPs and their associated cytochrome P450 reductases (CPRs), which transfer electrons from NADPH to their bound cytochrome P450^44,45^. To overcome this challenge, researchers have proposed that certain CYP–CPR systems, such as the one involved in dhurrin biosynthesis^46^, naturally assemble into membrane-bound nanoscale clusters termed “metabolons,” which facilitate both efficient electron transfer and substrate proximity channeling.

Inspired by these natural membrane-bound metabolic clusters, we investigated whether recruiting a cytochrome P450 (CYP) and its CPR into ER-targeted sMACs could mimic natural metabolons and improve pathway activity by promoting more efficient electron transfer between CYP and CPR (Fig 3a). To this end, we applied the multiPpID building block incorporating three PpIDs (ALFA nanobody, Crk SH3, and PTB), Dbp1n, and the ER-targeting motif Ubc6p(232–250) (Supplementary Note 1) as the basic unit for the ER-associated condensate. Crk SH3 and PTB domains were used to recruit CYP and CPR, respectively, forming the basis of ER-associated CYP/CPR synthetic metabolons.

**Figure 3.**
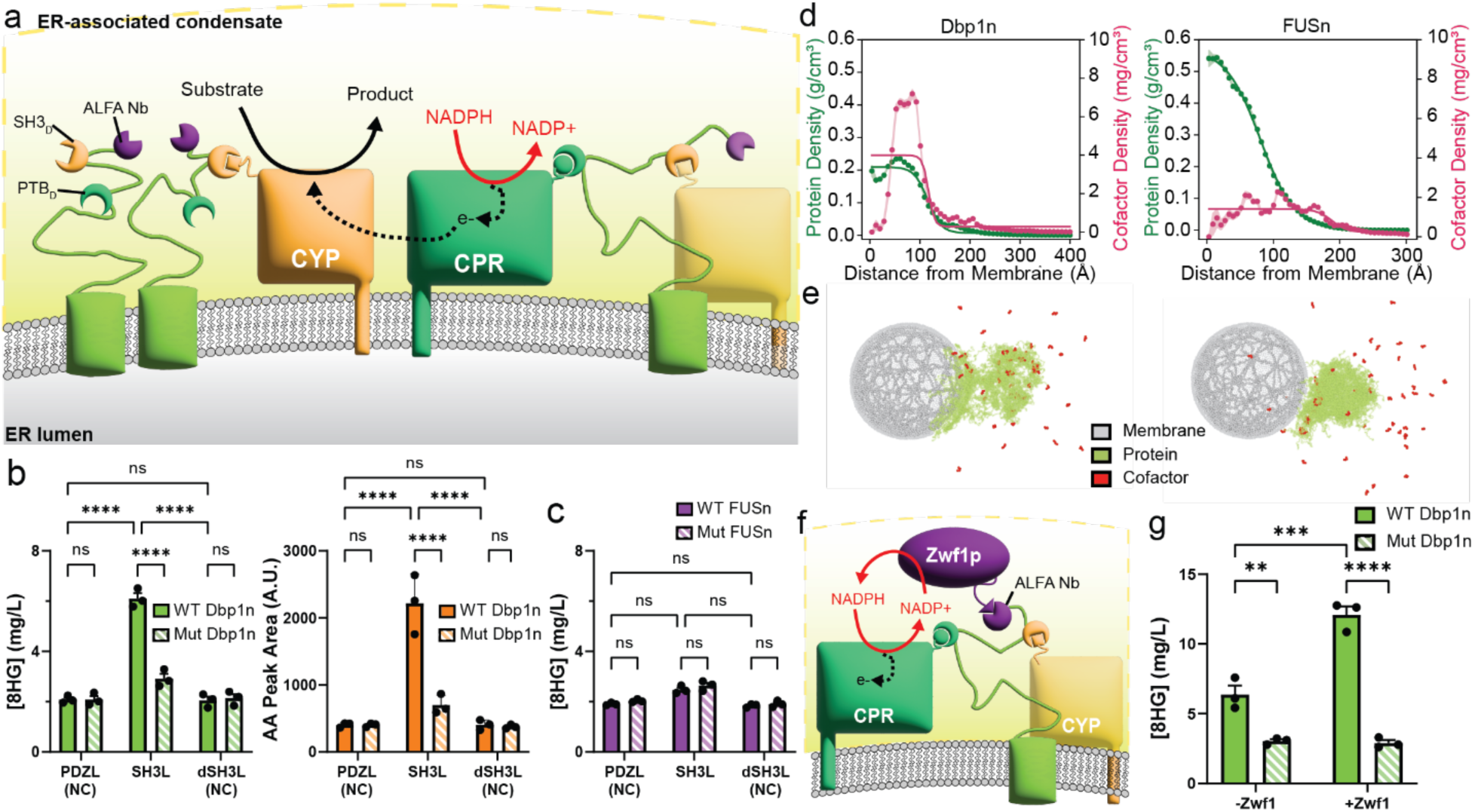
ER-associated condensate enhances cytochrome P450 productivity by co-localizing CYP and CPR along with elevated concentrations of NADPH. **(a)** Schematics of artificial metabolons for CYP-CPR systems. Recruitment of CYP and CPR into sMACs is mediated by PpIDs, promoting more efficient electron transfer. **(b)** Titers of CYP products (8HG in green and AA in orange) were significantly higher from fermentations using strains with recruitment of CYP and CPR into ER sMACs made of Dbp1n compared to control strains without recruitment (“NC”) or where sMACs fail to form by substituting Mut Dbp1n in place of WT Dbp1n. **(c)** The increase in 8HG titers was not observed when FUSn was used to construct ER sMACs, suggesting effects beyond simple substrate channeling. **(d)** Radial protein and NADPH density profile of simulated sMAC in coarse-grained direct coexistence molecular dynamics simulation. Points represent the time-averaged density over equilibrated simulation frames at a given radial position, shaded region represents the standard error, and curve represents a generalized logistic fit to data. **(e)** Representative coarse-grained molecular dynamics frame where the surface of a membrane-bound organelle (red) contains membrane-tethered IDRs (green) that recruit cytosolic IDRs (blue) and NADPH (pink). The leftmost frame represents recruitment where Dbp1n is the membrane-tethered and cytosolic IDR, while the right depicts a frame where FUSn is the IDR. **(f)** Schematic of sMAC-localized NADPH-regeneration cycle upon recruitment of *Sc*Zwf1p along with CYP and CPR into the artificial metabolon. **(g)** 8HG titers increased approx. two-fold upon incorporation of *Sc*Zwf1p into the CYP-CPR artificial metabolon and about six-fold higher than either no recruitment of enzymes into ER sMACs or no formation of sMACs where Mut Dbp1n was used. Two-way ANOVA was used to compare the sample data between each group of three biological replicates. All data are shown as the mean ± s.e.m. with the following significance notation: ns denotes not significant; ** denotes p<0.01; *** denotes p<0.005; **** denotes p<0.001.

As proof of concept, we recruited two CYP-CPR systems into ER sMACs to create artificial metabolons: 1) geraniol 8-hydroxylase from *Catharanthus roseus* (*Cr*G8H) to produce 8-hydroxygeraniol (8HG) and 2) CYP71AV1 from *Artemisia annua* to produce artemisinic acid (AA) (Supplementary Fig 2). We observed that the titers of 8HG and AA were approximately three-and five-folds higher, respectively, compared to controls where enzymes were not recruited to condensates (Fig 3b). These results suggest that the enhancement of CYP activity through sMAC application is generalizable across multiple CYP systems.

Interestingly, the increases in 8HG titers were only observed when Dbp1n was used as the IDR compared to FUSn (Fig 3c). We hypothesized that the higher product titers observed with Dbp1n resulted from enhanced NADPH partitioning into Dbp1n condensates relative to those formed by FUSn, as supported by both MD simulations and experimental data described above (Table 1). To determine whether this phenomenon extends to ER-associated sMACs, we modified our computational simulations to incorporate a lipid membrane model with anchored IDRs (Fig 3d and 3e and Supplementary Video 3 and 4, and “Direct coexistence simulation of membrane-tethered IDRs” in Methods). These simulations revealed strong NADPH partitioning into Dbp1n condensates (K_p_ ∼13), whereas only weak partitioning was observed with FUSn (K_p_ ∼1.4) (Table 1). Because NADPH is the electron donor required for CYP catalysis, its local availability can profoundly influence activity^47,48^. Thus, the elevated NADPH concentrations achieved through Dbp1n-mediated partitioning within sMACs are likely to enhance CYP function and, ultimately, increase product titers.

To further assess the impact of local NADPH concentration on CYP activity, we recruited *Sc*Zwf1p, a glucose-6-phosphate dehydrogenase that regenerates NADPH from NADP⁺, using the ALFA nanobody into ER sMACs (Fig 3f). This design enabled *in situ* recycling of the cofactor, establishing a closed-loop system to sustain CYP catalysis. Recruitment of *Sc*Zwf1p led to an approximately twofold increase in 8HG titers (12.1 mg/L vs 6.57 mg/L, Fig 3g), indicating that local NADPH regeneration further enhanced CYP-dependent production. Together, these results demonstrate that ER sMACs not only facilitate efficient electron transfer between CYP and CPR but also remodel the biochemical microenvironment, specifically by elevating and recycling NADPH, in ways that significantly enhance CYP activity.

### PM-associated synthetic transport metabolon enhances xylose uptake and utilization

To further develop sMACs as a versatile platform for organizing biological processes on membranes, we sought to extend it to the plasma membrane (PM). The PM not only serves as the physical boundary between the cell and its environment but also represents a critical interface for engineering access to non-native substrates. By positioning synthetic scaffolds at this boundary, it may be possible to couple transport with immediate downstream processing, thereby overcoming transport barriers that limit substrate utilization. A notable example is xylose utilization in engineered *S. cerevisiae* for lignocellulosic biomass conversion. Although yeast encodes a xylose reductase (XR) and a xylitol dehydrogenase (XDH), it cannot grow solely on xylose without extensive genetic modifications^49^. Inefficient uptake and poor integration of xylose into central metabolism have been hypothesized as major limitations. To address this, Thomik et al. developed an “artificial transporter metabolon”^50^, in which the galactose transporter Gal2p was brought to proximity of a xylose isomerase (XI) through structured protein scaffolds. Despite the enzyme’s slow kinetics (k_cat_ ∼5 s⁻¹; K_M_ for xylose ∼62 mM; k_cat_/K_M_ ∼0.080 s⁻¹ mM⁻¹)^51^, this design enhanced xylose uptake and conversion to ethanol over >150 hours under anaerobic conditions relative to non-scaffolded controls, an effect attributed to suppression of xylitol inhibition on XI in the artificial metabolon^50^.

Building on this concept, we next investigated whether PM-associated sMACs could facilitate xylose uptake and assimilation. We applied the multiPpID building block with the three PpIDs (ALFA nanobody, Crk SH3, and PTB), Dbp1n, and a plasma membrane-targeting motif Gap1p(550–602) as the basic unit of the PM sMACs (Fig 4a, Supplementary Note 1). The designed cargo proteins included a hexose transporter (HXT, i.e. Hxt7p), a XR from *Scheffersomyces stipitis* (formerly *Pichia stipitis*), i.e. *Ps*Xyl1, and a XDH from the same organism i.e. *Ps*Xyl2. We reasoned that scaffolding XR and XDH at the PM would establish an artificial transporter metabolon that not only bypasses the xylitol inhibition and kinetic limitations of XI but also exploits the higher catalytic efficiencies of XR (k_cat_ ∼7 s⁻¹; K_M_ for xylose ∼42 mM; k_cat_/ K_M_ ∼0.17 s⁻¹ mM⁻¹)^52^ and XDH (k_cat_ ∼17 s⁻¹; K_M_ for xylitol ∼26 mM; k_cat_/K_M_ ∼0.67 s⁻¹ mM⁻¹)^53^ relative to XI. Because both enzymes are tightly regulated by cofactor availability (NADH/NAD⁺ and NADPH/NADP⁺), we hypothesized that sMAC-mediated enrichment of cofactors would further enhance flux. In particular, recruiting an engineered NADH-dependent XR^54^ together with a NAD⁺-dependent XDH should create a localized NADH/NAD⁺ regeneration cycle, analogous to the NADPH recycling enabled by *Sc*Zwf1p in ER sMACs, thereby accelerating conversion of xylose to xylulose, and ultimately improving assimilation (Fig 4a).

**Figure 4.**
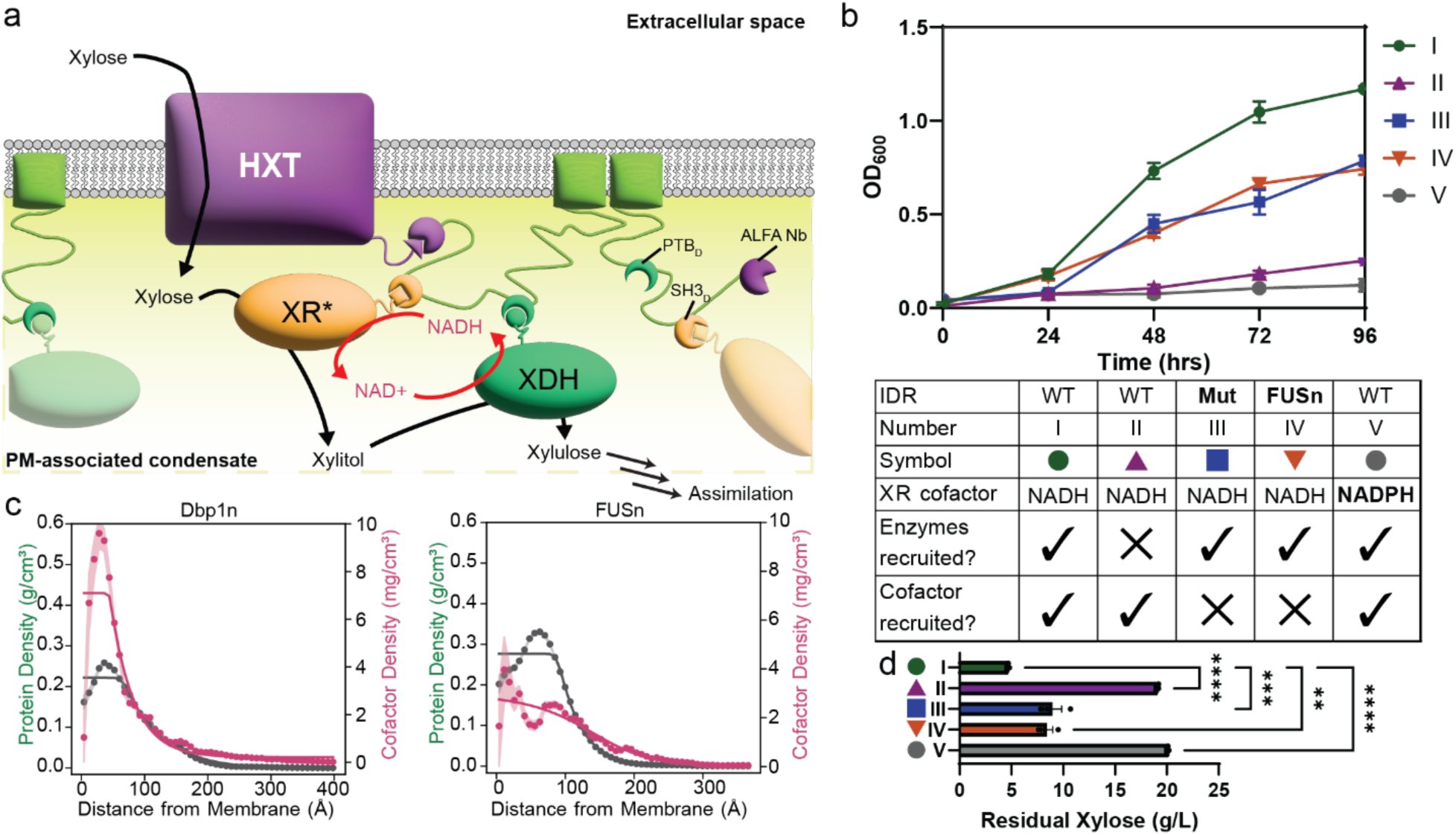
PM-associated condensate promotes xylose uptake and assimilation by localizing co-factor regeneration. **(a)** Schematic of PM sMACs recruiting a transporter (HXT), xylose reductase (XR, *Ps*Xyl1), and xylitol dehydrogenase (XDH, *Ps*Xyl2), creating a xylose uptake metabolon driven by *in situ* NADH/NAD^+^ regeneration cycle. **(b)** Time course of engineered yeast strains growing on xylose medium showed that in the absence of enzyme recruitment (II), condensate formation (III), cofactor recruitment (IV), or cofactor recycling (V), a growth defect relative to strains forming the complete transport metabolon (I) was observed. Error bars represent standard error of three biological replicates. **(c)** Radial protein and NADH density profile of simulated sMAC in coarse-grained direct coexistence molecular dynamics simulation. Points represent the time-averaged density over equilibrated simulation frames at a given radial position, shaded region represents the standard error, and curve represents a generalized logistic fit to data. **(d)** The strain with condensate formation, enzyme recruitment, cofactor recruitment, and cofactor recycling consumed a significantly higher amount of xylose from growth medium relative to all control strains, suggesting that PM sMACs could create more optimal local biochemical environments to enhance uptake and assimilation. One-way ANOVA was used to compare the sample data between each group of three biological replicates. All data are shown as the mean ± s.e.m. with the following significance notation: ** denotes p<0.01; *** denotes p<0.005; **** denotes p<0.001.

To test this hypothesis, we constructed five configurations of the artificial transporter metabolon (Fig 4b, I–V) and introduced each into a yeast strain lacking all hexose transporters (EBY.VW4000, a gift from Eckhard Boles). When cultured on xylose, only strains in which NADH-dependent XR and NAD⁺-dependent XDH were recruited to the artificial metabolon exhibited growth, irrespective of the IDR variant used (Fig 4b). In contrast, no growth was observed in control strains where XR and XDH were not recruited into PM sMACs (II) or when XR was replaced by a NADPH-dependent variant (V). These results indicate that both xylose channeling, as demonstrated by Thomik et al. ^50^, and cofactor recycling between scaffolded XR and XDH (NADH/NAD⁺ regeneration) are essential for promoting flux through the localized pathway.

We additionally observed a further increase in growth when Dbp1n was used as the IDR to promote sMAC formation (I), compared with either the non-condensing mutant variant (III) or FUSn (IV). This improvement is consistent with the enhanced NADH partitioning into Dbp1n condensates we observed (Table 1), analogous to the NADPH enrichment observed in ER sMACs. Also supporting this mechanism, MD simulations incorporating a lipid membrane model revealed strong NADH partitioning into membrane-associated sMACs (K_p_ ∼19.5) (Table 1 and Fig 4c). Experimental results further corroborated these findings: the strain expressing Dbp1n PM sMACs (I) consumed the most xylose of all strains, followed by strains with localized NADH/NAD⁺ regeneration (III and IV), which consumed more than half of the initial xylose in the growth medium (Fig 4d). Taken together, these results indicate that recruitment of XR and XDH into PM sMACs enhances xylose uptake and assimilation by establishing a favorable local biochemical environment that supports efficient cofactor recycling.

## Discussion

Our study establishes synthetic Membrane-Associated Condensates (sMACs) as a programmable platform for organizing and controlling metabolic reactions through the emergent properties of phase separation at intracellular membranes. Unlike conventional scaffolding strategies in synthetic biology that have typically focused on colocalizing enzymes to improve reaction efficiency^55,56^, sMACs can simultaneously tune enzyme recruitment, cofactor availability, and potentially small molecule metabolite concentration. These mechanisms collectively expand the design space for metabolic engineering.

In this work, we establish a quantitative framework linking IDR sequence to redox cofactor partitioning, enabling rational design of sMACs for enhancing specific metabolic functions. Using coarse-grained molecular dynamics simulations, validated by experimental partitioning assays, we demonstrated that Dbp1n selectively enriches NADH and NADPH within condensates, in contrast to FUSn. These results connect sequence-level motifs to emergent biochemical environments, showing that the capacity of condensates to influence metabolism extends beyond protein recruitment to include control over cofactor concentrations^27,57^. Importantly, the agreement between simulations and experiments suggests that computational models can serve as predictive tools for designing condensates with tailored partitioning behaviors^58^.

The case studies presented here demonstrate the benefits of cofactor partitioning and recycling within the synthetic condensates. In ER-associated sMACs, Dbp1n-mediated enrichment of NADPH enhanced CYP activity by improving electron transfer efficiency between CYP and CPR. This effect was directly tied to increased product titers, establishing that condensates can influence catalytic performance by altering the redox state of their microenvironment. Similarly, in PM-associated sMACs, NADH enrichment supported the catalytic conversion of xylose to xylulose by XR and XDH, enabling more efficient xylose uptake and assimilation. When combined with localized cofactor recycling by recruiting *Sc*Zwf1p to regenerate NADPH within ER sMACs, or pairing NADH-dependent XR with NAD⁺-dependent XDH in PM sMACs, these condensates establish localized redox cycles that reduce diffusive losses, enhance electron transfer efficiency, and ultimately drive flux beyond what is achievable in the bulk cytosolic milieu. In both ER and PM systems, enhancement in metabolic activities could not be explained by enzyme colocalization alone, but required the additional improvement afforded by cofactor partitioning and recycling. These results illustrate how condensates function as “microreactors” that operate as “synthetic organelles” where enzyme recruitment and cofactor availability can enhance pathway fluxes^47,48^.

Our work extends the concept of the metabolon by constructing a synthetic analogue that leverages membrane anchoring and phase separation to create distinct biochemical environments. Whereas natural metabolons and earlier synthetic scaffolds primarily rely on protein–protein interactions in the cytosol^35,56,59^, sMACs demonstrate that phase-separated assemblies on membranes can provide additional benefits and layers of control. These artificial assemblies are thus conceptually analogous to lipid rafts, where selective partitioning of components into dynamic membrane microdomains is thought to regulate local signaling and metabolism^60^. By anchoring condensates to organelles such as the ER and plasma membrane, we show that sMACs can impose spatial organization of metabolic enzymes and transporters, with the ability to tune redox availability at defined subcellular locations. This modularity suggests that sMACs could, in principle, be adapted to reprogram flux across diverse pathways and compartments in which transport and cofactor balance are limiting^50,61,62^.

Despite this versatility, not all metabolic pathways are likely to benefit uniformly from sMACs. This is reflected in the different fold improvements observed for the two CYP–CPR systems tested. Recruitment into ER-sMACs increased 8HG production by approximately three-fold but enhanced AA titers by nearly five-fold. Such variations likely stem from intrinsic properties of the enzymes themselves, including differences in catalytic turnover, coupling efficiency, or binding affinity for NADPH, which may influence the extent to which local cofactor enrichment enhances activity. Another key limitation of the current system is that, although we have demonstrated selective enrichment of enzymes and cofactors using appropriate IDRs, the complementary requirement, i.e. active exclusion of inhibitory or competing molecules, has not yet been achieved. In cases where undesired metabolites, inhibitors, or cross-reactive substrates diffuse freely into the condensate, pathway performance may remain constrained.

Potential solutions include situating sMACs within a membrane-bound organelle to combine the selectivity of lipid bilayers with the concentrating effects of condensates, or engineering multi-layered condensate architectures in which an outer shell consumes, buffers, or otherwise shields the core from undesirable components. Such strategies would expand the range of pathways that can be effectively controlled using sMACs and are actively being explored.

Future work could further explore how condensates influence both their recruited proteins and the membranes on which they assemble. While our study focused primarily on cofactor partitioning, the altered biochemical environment of sMACs likely also impacts protein conformation^63^ and enzymatic activity^64^, potentially enhancing or impairing pathway performance in ways that were only implicitly observed here. Accordingly, a robust methodology for tuning condensate properties to match the biochemical requirements of recruited enzymes will be essential. Such an approach could combine predictive computational modeling with experimental high-throughput screening to systematically optimize features such as partitioning^58^, pH buffering^64^, or charge. Equally important is the understudied interplay between condensates and membranes: membrane-associated assemblies could impose mechanical stress^65^, perturb trafficking^65^, or alter lipid homeostasis^66^, with consequences for cellular fitness and metabolism. Additionally, while small molecule partitioning is an equilibrium property, the cell is inherently a nonequilibrium system. While our results suggest that Dbp1n sMACs can reduce diffusive losses of redox cofactors, additional approaches are needed to investigate the impact of condensate viscoelastic properties on reaction dynamics and vice-versa under nonequilibrium conditions. Deeper biophysical and cellular studies will thus be critical for designing next-generation sMACs that integrate cofactor recycling with precise control over enzyme activity while maintaining membrane compatibility.

Beyond metabolic engineering, these findings shed light on the fundamental mechanisms by which cells use phase separation to regulate biochemical processes. Our results parallel emerging evidence that native membrane-associated condensates influence ion transport^16^, cofactor partitioning^23^, and energy metabolism^8,36^. By building synthetic mimics, we not only replicate these functions but also extend them in programmable ways in living cells. Future work will focus on expanding the repertoire of cofactors and metabolites whose partitioning into synthetic condensates can be tuned, and on integrating computational design with high-throughput experimental screens. Such efforts could yield predictive assembly of condensates in targeted subcellular compartments optimized for diverse biotechnological applications, from biofuel production to pharmaceutical biosynthesis.

## Acknowledgements

We thank Professor Eckhard Boles at Goethe University for his generous gift of hexose transporters knockout strain EBY.VW4000 used as the base strain for the xylose uptake experiments. We also thank previous members of the Avalos lab Tess Kichuk and Lisset Duran for providing plasmids expressing ER and mitochondrial markers pTK11 and pLD67, respectively. The training and support provided by Dr. Gary Laevsky and Dr. Sha Wang of the Confocal Imaging Facility (CIF) in the Department of Molecular Biology for the microscopy experiments are greatly appreciated. Molecular dynamics simulations were performed using the Princeton Research Computing resources at Princeton University.

This work was supported by the Princeton Biomolecular Condensate Program, the Air Force Office of Scientific Research (AFOSR) Multidisciplinary University Research Initiative (MURI, FA9550-20-1-0241 to J.L.A.) and partially supported via National Science Foundation (NSF) through the Princeton University (PCCM) Materials Research Science and Engineering Center (DMR-2011750) to J.A.J. V.J. is supported by NSF GRFP (DGE-2039656) and a Gordon Wu Fellowship. M.T.W. was supported by the NSF GRFP (DGE-1656466). We thank members of the Avalos, Brangwynne, and Joseph labs for valuable discussions and feedback throughout this project.

## Author contributions

K.S., V.J., and J.L.A. designed the research. K.S., and E.M.G. conducted experiments for characterizing sMAC-expressing yeast strains and xylose uptake and assimilation. K.S. additionally conducted experiments for 8HG and AA biosynthesis. V.J. conducted computational simulations and experimental validation for cofactor partitioning. M.T.W. and K.S. designed the PpID system. K.S., V.J., and J.L.A. wrote the manuscript and created the figures. All authors revised the text and figures.

## Methods

### Strains and growth media

The base *S. cerevisiae* strain for all experiments except xylose assimilation experiments was CEN.PK2-1C (MATa ura3-52 trp1-289 leu2-3,112 his3Δ1 MAL2-8C SUC2). Yeast cultures were grown in YPD medium (10 g L^−1^ Bacto yeast extract, 20 g L^−1^ Bacto peptone, 20 g L^−1^ glucose). Selection of markers (i.e. URA3) was performed in synthetic complete (SC) medium (1.5 g L^−1^ yeast nitrogen base without amino acids or ammonium sulfate, 5 g L^−1^ ammonium sulfate, 36 mg L^−1^ inositol, and 2 g L^−1^ amino acid drop out mixture; 20 g L^−1^ glucose).

The base *S. Cerevisiae* strain used for xylose uptake and assimilation experiments was EBY.VW4000 (MATa ura3-52 trp1-289 leu2-3,112 his3Δ1 MAL2-8C SUC2 Δhxt1-17 Δgal2 Δstl1::loxP Δagt1::loxP Δmph2::loxP Δmph3::loxP, a gift from Eckhard Boles, Goethe University)^67^. Because all hexose transporters are knocked out in EBY.VW4000, cultures were grown in YPM media (10 g L^−1^ Bacto yeast extract, 20 g L^−1^ Bacto peptone, 20 g L^−1^ maltose), with maltose taking the place of glucose. Selection of markers (i.e. URA3) was performed in synthetic complete (SC) medium (1.5 g L^−1^ yeast nitrogen base without amino acids or ammonium sulfate, 5 g L^−1^ ammonium sulfate, 36 mg L^−1^ inositol, and 2 g L^−1^ amino acid drop out mixture; 20 g L^−1^ glucose)

GoldenGate assembly reactions were transformed into chemically competent *Escherichia coli* prepared from strain NEB® Turbo (NEB). Transformed cells were selected on LB containing the corresponding antibiotics chloramphenicol, ampicillin or kanamycin.

### Plasmids and yeast strains construction

Yeast expression vectors were built using GoldenGate assembly as described in the Yeast Toolkit (YTK) system^68^. All plasmids used in the study are listed in Supplementary Table 2, sorted by Figures in which they were used.

The required yeast expression cassettes are transformed into the base strains as described for the specific sets of experiments below. CRISPR-assisted chromosomal integration into the yeast genome was achieved by cotransforming a CEN6/ARS4 CRISPR plasmid (containing Cas9, a guide RNA for targeting the appropriate locus and a URA3 auxotrophic marker) with NotI-linearized repair DNA designed to integrate the gene(s) of interest. Cells were plated on uracil-dropout medium. Correct chromosomal integration was confirmed by PCR, and curing of CRISPR plasmid was completed by outgrowth in non-selective medium followed by plating onto 5FOA supplemented (1 g L^−1^ 5FOA (Zymo Research)) medium (a colony will only grow on 5FOA-supplemented medium if the CRISPR plasmid has been removed). All transformations were performed using a standard lithium acetate method, and cells were plated onto selective SC agar plates with 2% glucose or maltose (for CEN.PK2-1C or EBY.VW4000, respectively). Strains used for constructing the experimental strains are listed in Supplementary Table 3. Individual colonies (biological replicates) were picked directly from this transformation plate and grown independently for further analysis.

### Confocal fluorescence microscopy and analysis

For confirmation of protein localization and sMACs formation, strains were grown to saturation in SC medium with 2% glucose, diluted 50-fold into fresh medium and grown for a further 4–8 h. Cultures were then washed and resuspended in 1× PBS before adhering to 96-well glass bottom plates (Cellvis) pretreated with concanavalin A for imaging. Pretreatment of 96-well plates were done by pipetting 15 mL of 1 mg mL^−1^ concanavalin A (ConA) in 20 mM NaOAc and centrifuged for 1 min at 1,000 RPM. Plates were then incubated at room temperature for 1 h. Following incubation, wells were washed twice with 100 mL phosphate-buffered saline (PBS) and allowed to dry for 1 h at room temperature, after removing most of the last wash volume. Once the wells were dry, 50 mL of the resuspended cells (see above) were added to each well and the plate was centrifuged for 2 min at 1,000 RPM.

Live yeast imaging was performed on a Nikon W1 SoRa, controlled via Nikon Elements 6.01, at room temperature using a 63× oil objective (NA = 1.49) with 4× magnification for a pixel size of 0.027 μm. Yeast cells were imaged in 3D (35–50 z sections, 0.2 μm apart). The mTagBFP2, Venus, and mRuby2 fluorophores were excited with 405 nm, 488 nm, and 561 nm excitation respectively. Images were denoised using Nikon Elements and then processed in ImageJ.

### Measurement of NADH/NADPH partitioning

Overnight cultures of yKS331 (pGAL-utTrimer-ScDbp1n (2-214)-mCherry-Cvk_SH3-PTB-tEno1) were grown in SC-Leu for 16 hours, then expression was induced after diluting the overnight 1:1000 in SC-Leu with 0.2% glucose and 1.8% galactose. Yeast cells were grown for 48 hours at 30°C, then harvested by centrifugation at 7,500 x g for 20 minutes at 4°C. Cell pellets were resuspended in 10 mL L^−1^ Tris-buffered saline-TBS, pH 7.6, and flash-frozen. Cells were lysed using a CryoMill system and combined with 20 mM Tris, 150 mM NaCl buffer supplemented with 1 mM PMSF, 2 mg mL^−1^ of DNase and 100 µL of commercial protease inhibitor cocktail.

Cell lysate was centrifuged at variable salt concentrations and temperatures to induce movement of the target construct from the dense to dilute phase, as described in Li et al^69^. Salt concentrations and temperatures were chosen based on the phase diagrams of Dbp1n at 150 mM NaCl and 1.5M NaCl. The pellet was washed with 20 mM Tris, 1M NaCl and centrifuged again at 25°C, then washed again with 20 mM Tris, 1M NaCl, 3M urea and centrifuged again at 25°, then heated to 60°C and centrifuged again. Each time, the supernatant was saved, filter sterilized, and the mCherry fluorescence (excitation 587 nm, emission 610 nm) was measured with a Tecan Infinite 200 Pro F-plex. The third supernatant had the highest fluorescence/A280 ratio and so was desalted and concentrated using a 30 KDa Amicon Ultra-15 Centrifugal Filter Unit. Proteins were flash-frozen in liquid nitrogen and stored at −20°C.

To verify droplet formation, protein solutions were diluted to 20 µM in 20 mM Tris buffer. Solutions were left to sediment at 4°C for 10 minutes, centrifuged at 2000 RPM for 5 minutes to settle droplets, and imaged using a Nikon TI-2 Eclipse widefield microscope under 100x oil immersion lens.

To quantify small molecule partitioning, 20 µM protein solution was combined with 100 µM of either NADPH or NADH in Tris buffer, pH 7.6. Solutions were left to sediment at 4°C for 10 minutes, then centrifuged at 12400 RPM for 1 hour. The dense and dilute phases were mechanically separated and resuspended to 50 µL working volume with 20 mM Tris. NADH or NADPH concentration was measured through fluorescence (excitation 340 nm, emission 460 nm) compared to a standard curve, and protein concentration was measured through absorbance at 280 nm and mCherry fluorescence. Partition coefficients were determined by taking the ratio of dense and dilute phase concentrations across three replicates.

### 8-Hydroxygeraniol (8HG) fermentation

#### Strain development

Starting with the parental strain CEN.PK2-1C, we deleted OYE2 and OYE3 ORFs using homologous recombination and selection with nourseothricin and hygromycin markers, respectively, to create yMW941. These genes were knocked-out to eliminate their known activities on geraniol and 8HG that lead to loss of 8HG as “shunt products”^70^. A set of 4 genes, acetoacetyl-CoA thiolase/hydroxy-methyl-glutaryl-CoA reductase from Enterococcus faecalis (*Ef*MvaE), Isopentenyl-Diphosphate Delta Isomerase (*Sc*IDI1) and a mutated Farnesyl pyrophosphate synthetase (*Sc*Erg20(F96W/N144W)) from *S. cerevisiae*, and Geraniol Synthase from *Ocimum basilicum* (*Ob*GES), was made into a markerless multigene integration plasmid^68^ (pKS557) with upstream homology of 461bp and downstream homology of 578 bp around the XI-1 locus^71^. To integrate these genes, pKS557 was linearized by NotI and introduced into yMW941 along with a custom CRISPR plasmid (pKS208) expressing both Cas9 and sgRNA targeting the XI-1 locus. Briefly, the CRISPR-assisted integration was directed by the sgRNA with spacer sequence (5’-GCAATGCGATGTTAGTTTAG) targeting Cas9 to the XI-1 locus, enabling markerless integration by the linearized multigene cassettes and creating the base geraniol-producing strain. To cure out the CRISPR/Cas9 plasmid targeting XI-1 (pKS208), the geraniol-producing strain was cultured overnight in YPD with 2% glucose and plated onto YPD agar plate supplemented with 1 g L^−1^ 5FOA to create yKS109. Different fusions of protein-protein interaction ligands (PDZL, SH3L, PTBL, and their inactive variants) and the cytochrome P450 geraniol 8-hydroxylase (*Cr*G8H) along with those of the cytochrome P450 reductase (*Cr*CPR) from *Catharanthus roseus* were made into GAL-inducible expression plasmids (pKS1337, 1003, and pKS1077) with homologous regions targeting XII-5 locus^71^. These plasmids were linearized by NotI and introduced into yKS109 via CRISPR-assisted integration by cotransforming with pKS214 to create strains yKS228 and yKS231. Plasmids containing gene cassettes expressing ER sMAC base unit (ALFA nanobody, SH3, and PTB PpIDs fused to Dbp1n, Mut Dbp1n, FUSn, or Mut FUSn) and *Sc*Ubc6p(233-250)) were made for integration into the X-4 locus as pKS1126, pKS1129, pKS1054, and pKS1057. These plasmids were linearized by NotI and introduced into yKS228 and yKS231 to create the final 8HG fermentation strains. All transformed strains were verified by colony PCR.

#### Sample preparation and test tube fermentation

Following yeast transformation, six freshly transformed colonies were randomly picked and verified for proper integration with PCR. Three positive isolates were grown overnight in 1 mL SC media with 2% (w/v) glucose at 30°C in 24-well plate. On the next day, the overnight culture was used to inoculate 3 mL of SC media with 0.2% (w/v) glucose and 1.8% (w/v) galactose in 14 mL test tube with snap cap (Greiner Bio-One) at a starting OD600 of 0.2 and incubated at 30°C for 72 hr with shaking at 265 rpm.

#### Analytical methods and analysis

To quantify 8HG production, 600 μL of the post-fermentation culture was sampled and mixed with 600 μL of ethyl acetate (Sigma Aldrich). The mixture was vortexed for 30 min to extract organic products from the culture, followed by centrifugation at 12,500 g for 30 min at 4°C to separate the organic and aqueous layers. 550μL of the resulting organic (top) layer was transferred to a new microcentrifuge tube and centrifuged at 12,500 g for 30 min at 4°C again to remove any residual aqueous phase and yeast cells. Then 500μL of the resulting extract was taken for GC-MS analysis. 8HG was measured using an Agilent 7890B GC System. For analysis, samples were injected without split (0.5μL injection volume) with a constant flow of 1.5 mL/min. Separation of samples was achieved using a DB-Wax column (30 m length, 0.25 mm diameter, 0.5 m film) and a gradient method. The inlet temperature was 250 °C. The oven temperature was set to 70 °C initially and maintained for 5 minutes. Subsequently, the temperature was increased at a rate of 20 °C/min until reaching 230 °C, and then held for 3 minutes. Quantification of samples was performed using MS targeted detection of 8HG (121 m/z). The retention time for 8HG was ∼13.0 min. The concentration of the analyte was determined by comparing the integrated peak areas with a 8HG standard purchased from Toronto Research Chemicals.

### Artemisinic acid (AA) fermentation

#### Strain development

Starting with the parental strain CEN.PK2-1C, we introduced a set of 4 genes, acetoacetyl-CoA thiolase/hydroxy-methyl-glutaryl-CoA reductase from *Enterococcus faecalis* (*Ef*MvaE), Isopentenyl-Diphosphate Delta Isomerase (*Sc*IDI1) and wild-type Farnesyl pyrophosphate synthetase (*Sc*Erg20) from *S. cerevisiae*, and Amorphadiene Synthase from *Artemisia annua* (*Aa*ADS) fused to a solubilization-enhancing domain GB1^72^, was made into a markerless multigene integration plasmid^68^ (pKS709) with homologies around the XI-1 locus^71^. To integrate these genes, pKS709 was linearized by NotI and introduced into CEN.PK2-1C along with a custom CRISPR plasmid (pKS208) in a similar manner to pKS557 for creating yKS109 described above to create the base amorphadiene-producing strain yKS122. The accessory downstream enzymes *Aa*ADH1 and *Aa*ALDH1 were then introduced by transformation of NotI-linearized pKS1304 into yKS122 and integrated into the LEU2 locus with a LEU2 marker to create yKS342 to ensure subsequent conversion of intermediates into AA in the final strains. Different fusions of protein-protein interaction ligands (PDZL, SH3L, PTBL, and their inactive variants) and the cytochrome P450 amorpha-4,11-diene 12-monooxygenase (*Aa*CYP71AV1) along with those of the cytochrome P450 reductase (*Aa*CPR) and cytochrome b5 (*Aa*CYB5) from *A. annua* were made into GAL-inducible expression plasmids (pKS1324, pKS1266, and pKS1269) with homologous regions targeting XII-5 locus^71^. These plasmids were linearized by NotI and introduced into yKS342 via CRISPR-assisted integration by cotransformating with pKS214 to create strains yKS407 and yKS409. Plasmids containing gene cassettes expressing ER sMAC base unit (pKS1126 and pKS1129) as described above were linearized by NotI and introduced into yKS407 and yKS409 to create the final AA biosynthesis strains. All transformed strains were verified by colony PCR.

#### Sample preparation and shake flask fermentation

Following yeast transformation, six freshly transformed colonies were randomly picked and verified for proper integration with PCR. Three positive isolates were grown overnight in 1 mL SC media with 2% (w/v) glucose at 30°C in 24-well plate. On the next day, the overnight culture was used to inoculate 20 mL of SC media with 0.2% (w/v) glucose and 1.8% (w/v) galactose in 250 mL shake flask at a starting OD600 of 0.1 and incubated at 30°C for 72 hr with shaking at 250 rpm.

#### Analytical methods and analysis

To quantify AA production, 600 μL of the post-fermentation culture was sampled and mixed with 600 μL of ethyl acetate (Sigma Aldrich). The mixture was vortexed for 30 min to extract organic products from the culture, followed by centrifugation at 12,500 g for 30 min at 4°C to separate the organic and aqueous layers. 550μL of the resulting organic (top) layer was transferred to a new microcentrifuge tube and centrifuged at 12,500 g for 30 min at 4°C again to remove any residual aqueous phase and yeast cells. Then 500μL of the resulting extract was taken for GC-MS analysis. AA was measured using an Agilent 7890B GC System. For analysis, samples were injected with 20:1 split (0.5μL injection volume) with a constant flow of 1.5 mL/min. Separation of samples was achieved using a DB-Wax column (30 m length, 0.25 mm diameter, 0.5 m film) and a gradient method. The inlet temperature was 250 °C. The oven temperature was set to 150 °C initially and maintained for 3 minutes. Subsequently, the temperature was increased at a rate of 5 °C/min until reaching 250 °C, and then held for 8 minutes. Quantification of samples was performed using MS targeted detection of AA (206 m/z). The retention time for AA was ∼30.3 min.

### Xylose uptake and assimilation

#### Strain development

We first made a multigene plasmid containing the pentose phosphate pathway enzymes Tkl1p, Tal1p, Rpi1p, and Rpe1 that are known to improve xylose assimilation in *S. cerevisiae*^73^ through GoldenGate Assembly (pKS803). This plasmid is linearized by NotI and cotransformed with pKS208 into XI-1 locus using CRISPR-assisted integration in a manner like yKS109 and yKS122 described above to create yKS134. The hexose transporter capable of importing xylose, Hxt7p, was fused to ALFA ligand (ALFAL) and introduced into yKS134 via CRISPR-assisted integration into X-2 locus by cotransformaion of pKS179 with NotI-linearized pKS1043 to create yKS221.

Different fusions of protein-protein interaction domains ligands (SH3L, PTBL, and their inactive variants) and xylose reductase (*Ps*Xyl1, either wild type NADPH-utilizing or R276H NADH-utilizing variant) along with those of the xylitol dehydrogenase (*Ps*Xyl2) and xylulose kinase (*Ps*Xyl3) from *Scheffersomyces stipites* were then made into expression plasmids (pKS1068, pKS1070, and pKS1075) with homologous regions targeting XII-5 locus. These plasmids were linearized by NotI and introduced into yKS221 via CRISPR-assisted integration by cotransformating with pKS214 to create strains yKS244, yKS245, and yKS329. Plasmids containing gene cassettes expressing PM sMAC base unit (ALFA nanobody, SH3, and PTB PpIDs fused to Dbp1n, Mut Dbp1n, or FUSn and ScGap1p(550-602)) were made for integration into the X-4 locus as pKS1262, pKS1265, and pKS1309. These plasmids were linearized by NotI and introduced into yKS244, yKS245, and yKS329 to create the final xylose assimilation strains. All transformed strains were verified by colony PCR.

#### Sample preparation and test tube fermentation

Following yeast transformation, six freshly transformed colonies were randomly picked and verified for proper integration with PCR. Three positive isolates were grown overnight in 1 mL SC media with 2% (w/v) maltose at 30°C in 24-well plate. On the next day, the overnight culture was used to inoculate 3 mL of SC media with 2% (w/v) xylose in 14 mL test tube with snap cap (Greiner Bio-One) at a starting OD600 of 0.2 and incubated at 30°C for 96hr with shaking at 265 rpm.

#### Analytical methods and analysis

Cell growth is measured by taking 100μL of culture at indicated time points and diluting ten-fold into 900μL 1xPBS (8 g L^−1^ NaCl, 200 mg L^−1^ KCl, 1.44 g L^−1^ Na_2_HPO_4_, 245 mg L^−1^ KH_2_PO_4_, pH 7.4) in clear bottom 24-well plate. OD_600_ of diluted samples were then measured using a monochromator-based plate reader (TECAN infinite200PRO MPlex), the arbitrary units of which are approximately ten-fold lower than those measured with a spectrophotometer with a path length of 1 cm.

To quantify xylose, 600 μL of the post-fermentation culture was sampled and centrifuged at 3000 g for 5 min at 4°C to separate cells from supernatant. 550μL of the resulting supernatant was transferred to a new microcentrifuge tube and centrifuged again at 12,500 g for 30 min at 4°C to remove any residual cell debris. Then 500μL of the resulting supernatant was taken for HPLC analysis. an Agilent 1260 Infinity instrument (Agilent Technologies) and an Aminex HPX-87H ion-exchange column (Bio-Rad, catalog # 1250140), eluted with 5 mM sulfuric acid mobile phase at 55 °C. Chemical concentrations were estimated from the peak areas of refractive index detector compared to standards of known concentration.

### Computational methods

#### Identification of IDRs

A library of 3878 primary sequences of ATP-dependent RNA helicases were identified by functional annotations on UniProtKB across all taxa. A deep learning method (based off of ALBATROSS^74^) was applied to this dataset to identify disordered domains within each protein, calculate sequence properties and predict ensemble properties, including radius of gyration, end-to-end distance, asphericity, and Flory scaling exponent, for all identified IDRs. Radius of gyration (R_g_) normalized by the radius of gyration of an Analytical Flory Random Coil of the same number of amino acids is a measure of chain compaction, where lower R_g_ represents a higher degree of self-association. R_g_ has been shown to have an inverse power law correlation with critical temperature^75^. Charge patterning is represented by a parameter κ, which ranges from 0 (well mixed) to 1 (fully segregated)^76^, and patterning has previously been shown to be indispensable for mediating interactions between protein IDRs and nucleic acids^77^. Fractional charge per residue (FCR) has also previously been shown to be significantly higher for RNA binding IDRs.

To identify motifs likely responsible for liquid-liquid phase separation behavior, sequences that were greater than three standard deviations away from the mean fraction of charged residues, as well as sequences shorter than 50 amino acids in length, were filtered out of the identified IDRs. STREME^78–80^ was used to identify conserved motifs (defined as repeating units of 3-30 residues long, enriched over random chance) by e-value (probability of occurrence) and prevalence in the dataset.

FINCHES^28,81^, a mean-field method for calculating IDR—IDR interactions was used calculate self-interacting regions of the Dbp1n sequence, which corresponded with identified motifs. Critical residues in the identified motifs were mutated to reduce the critical temperature of the mutant Dbp1n sequence. A graphical overview of this method is given in Extended Data 1.

To identify IDR motifs likely implicated in binding negatively charged small molecules with adenosine moieties, we reasoned that motifs that contribute to cofactor partitioning would not be enriched across all IDRs present in the human proteome. We compared the motifs identified from running the same STREME algorithm on the previously described human IDRome to those identified in this filtered dataset^74^.

#### Developing Mpipi cofactor parameters

The Mpipi forcefield is a residue-level coarse grained model that has been found to quantitatively predict phase behavior for biomolecules such as proteins^28^ or RNA^82^. To generate Mpipi--compatible parameters for NADH, NAD+, NADPH, NADP+, and xylose, the all-atom parameterization method as described by Emelianova et. al was used^58^. Briefly, the Mpipi parameterization scheme considers the linear combination of harmonic bonded energies as well as nonbonded electrostatic and nonpolar interactions. Each monomer in a biopolymer is treated as an individual bead. Separate beads were parameterized to represent the adenosine monophosphate ribose, pyrophosphate, nicotinamide, and xylose. The nonpolar interactions are described using the Wang--Frenkel potential, where ε_i,j_ represents the strength of interaction between particles i and j, σ_i,j_ represents the Van der Waals radii, and ν_i,j_ and μ_i,j_ are parameters describing the decay of the interaction energy over distance.

To determine nonpolar interaction parameters, potential of mean force (PMF) calculations were performed using all-atom, explicit solvent representations of each amino acid and the functional groups of each small molecule. Umbrella sampling was used to sample energetically unfavorable states and the Weighted Histogram Analysis Method (WHAM) to remove the biased potential and recover the true probability distribution^83^. All-atom small molecule parameters were generated using Gaussian and Gaff2^84–86^. Simulations were run in GROMACS 2023.3 ^87^ with the IDP-specific forcefield AMBER ff99SBws to parameterize amino acids; ions and waters were parameterized with TIP4P/2005^88^. Each amino acid and small molecule pair was restrained in the x- and z- direction with a positional restraint of 1 J/mol-pm^2^ on every heavy atom while being pulled in the y-direction with force 6 J/mol-pm^2^, simulated across 50 windows between 0.1 and 2.0 nm center-of-mass distance. Hydrogen atoms were constrained using LINCS^89^; electrostatics were computed with particle-mesh Ewald summation, using a Coulombic cutoff of 1 nm^90^. Systems were neutralized and then sodium chloride was added to a final concentration of 150 mM. Wang--Frenkel^91^ ε values were calculated based on well depth and area from PMFs averaged across replicates from 3 independent thermal seeds while Wang--Frenkel σ values were calculated from Van der Waals volume^92^. Charged interactions are described using a Coulombic potential, where pyrophosphate beads contained a net −1 charge and nicotinamide beads in NADH and NADPH carried a net −1 charge. Ideal bond lengths were calculated from determining the geometric center for each bead from a crystal structure of each cofactor. Ideal angles were calculated similarly by measuring the angles between the geometric center of each bead mapped onto each cofactor’s crystal structure. Force constants were varied iteratively until the distribution of lengths and angles visited in coarse-grained simulation corresponded with that from all-atom simulation.

The parameters were validated by computing contact frequencies between a single-copy Dbp1n protein and small molecules above the saturation concentration in Mpipi coarse-grained simulations and comparing the resulting distribution to GROMACS all-atom simulations using REST2^93^ as an enhanced sampling method. This method was first validated by running on an ATP and FUS PLD system and compared with published results^94^. For the REST2 all-atom simulations, the starting configuration of the protein was generated using the ColabFold implementation of the AlphaFold 2 modeling pipeline^95,96^. The system was equilibrated for 50 ns each in NVT/NPT simulation using the ff99SBws forcefield with GAFF2 parameters for the ligand, using the Berendsen thermostat at target temperature of 300K and target pressure 1 bar in GROMACS. Bond angles and bond lengths were constrained with the LINCS algorithm^89^; electrostatics were calculated using the PME algorithm^90^. Six collective variables were used according to the protocol in Zhu et.al^97^: motif φ/Ψ angles were set to ideal left-handed helix values, right-handed helix values, beta sheet values, and polyproline values; radius of gyration based on Cα position; and backbone hydrogen bonding distance. The equilibrated protein-ligand structure was simulated in NVT using a velocity-rescaling algorithm^98^ for a 16-replica temperature ladder between 300-500K, with a replica attempted every 80 ps. Each replica was simulated for 0.5 µs, for a total of 8 µs. All analysis was carried out using MDTraj^99^ and MDAnalysis^100,101^.. The initial configuration for the coarse-grained simulations was built from the Cα coordinates in the all-atom simulation, then minimized, equilibrated, and simulated in the NVT ensemble using Langevin integrator at 300K until the number of contacts in every frame stabilized (at least 0.1 µs). All simulations were run in cubic boxes of box length 10 nm; the molar ratio of small molecules to protein was 32:1.

#### Direct coexistence simulation of cytosolic IDRs and small molecules

All coarse-grained simulations are conducted using LAMMPS (version Aug 29 2024)^102^. Simulation rendering is performed using OVITO^103^. A direct coexistence simulation is when the dense and dilute phases of the condensate are simulated simultaneously in the same system. For cytosolic condensates, slab geometries representing an axial slice of a spherical condensate are used to minimize finite-size errors. Interactions are modeled with the Mpipi force field^104^. To set up direct coexistence slab simulations, several copies of the desired protein are compressed in the NPT ensemble using the Langevin thermostat (T=150K, τ_T_ = 10 ps) and the Berendsen barostat (P=700 atm, τ_P_ = 10 ps). The simulation box is then expanded in one dimension to a slab for a total protein density of 0.1gm/cm^3^. NVT simulation is performed on the slab for 200 ns of equilibration and 500 ns production. For simulations with small molecules, parameterized cofactors are inserted into the simulation box at an equimolar ratio to protein molar concentration prior to equilibration and production runs. Dense and dilute phase boundaries are determined based off calculating the stationary points and inflection points of a super-Gaussian fit to the block-averaged density profile along the expanded axis of the slab. Critical temperatures are calculated using the law of coexistence densities and critical densities are calculated using the law of rectilinear diameters^28^. Small molecule partition coefficients are determined by the ratio of the density of small molecules within the protein dense phase to the density of small molecules within the protein dilute phase.

#### Direct coexistence simulation of membrane-tethered IDRs

A spherical membrane-bound organelle was built using a mesh scaffold as described in Fu et.al^105^. Parameters for the phospholipid membrane were adapted from the LAMMPS fluid membrane pair potential, scaled such that each phospholipid bead had diameter 5.2 nm, consistent with a previously determined thickness of the lipid bilayer^106^. Electrostatic fluid membrane parameters were modified to account for screening from implicit solvent. The scaffold network, transmembrane anchors and phospholipids were treated such that the only pairwise interactions between the IDP or small molecule beads were excluded volume interactions, described with a Lennard-Jones pair potential. The membrane system (comprised of phospholipid, scaffold, a transmembrane anchors) was minimized using the fire algorithm, then equilibrated for 50 ns in the NVT ensemble at 300K with the Langevin thermostat, then equilibrated for 50 ns in the NPT ensemble at 1 atm with the Berensden barostat. Flexible protein chains were covalently bonded to transmembrane anchor proteins with a harmonic bonded potential to represent membrane-tethered IDPs. Membrane-localized proteins and the membrane were again equilibrated in NVT/NPT. A compressed sphere of cytosolic proteins, prepared as described above, was combined with the membrane-tethered system, and the entire system was run for 50 ns equilibration and 200 ns production. Small molecules were added to the system similarly as described above in the cytosolic case.

Dense and dilute phase boundaries were found through fitting a generalized logistic curve to the block-averaged radial protein density profile outward from the surface of the spherical membrane-bound organelle, then finding the stationary and inflection point of the logistic curve. The small molecule partitioning coefficient is again determined by the ratio of the small molecule concentration in the protein dense phase to the small molecule concentration in the protein dilute phase.

#### Simulations at elevated salt concentration

A similar all-atom simulation protocol was done to determine small molecule parameters at 1.5M and 3M NaCl concentration as above using PMFs; atomistic salt concentrations were increased to either 1.5M or 3M. Derived parameters were combined with protein parameters at elevated salt concentrations as previously described^31^. Coarse-grained direct coexistence simulations were run as described above to determine the critical point of the system and small molecule partitioning.

### Statistics and Reproducibility

Statistical analyses were performed as described in the Figure legends using GraphPad Prism

10.4 (Dotmatics). All microscopy images presented were representative of at least two independent biological replicates.

## Data Availability

Plasmids generated in this study are available upon reasonable request. Sample MD simulation trajectories and scripts to set up and run simulations in LAMMPS have been deposited on Zenodo.

## Code Availability

We conducted all simulations in the open-source LAMMPS MD simulation engine (Updated 22 Aug 2024). Simulation setup and production scripts are available on Zenodo. Tutorials for running simulations are available on <u>GitHub</u>.

## Extended Data

**Extended Data Figure 1.**
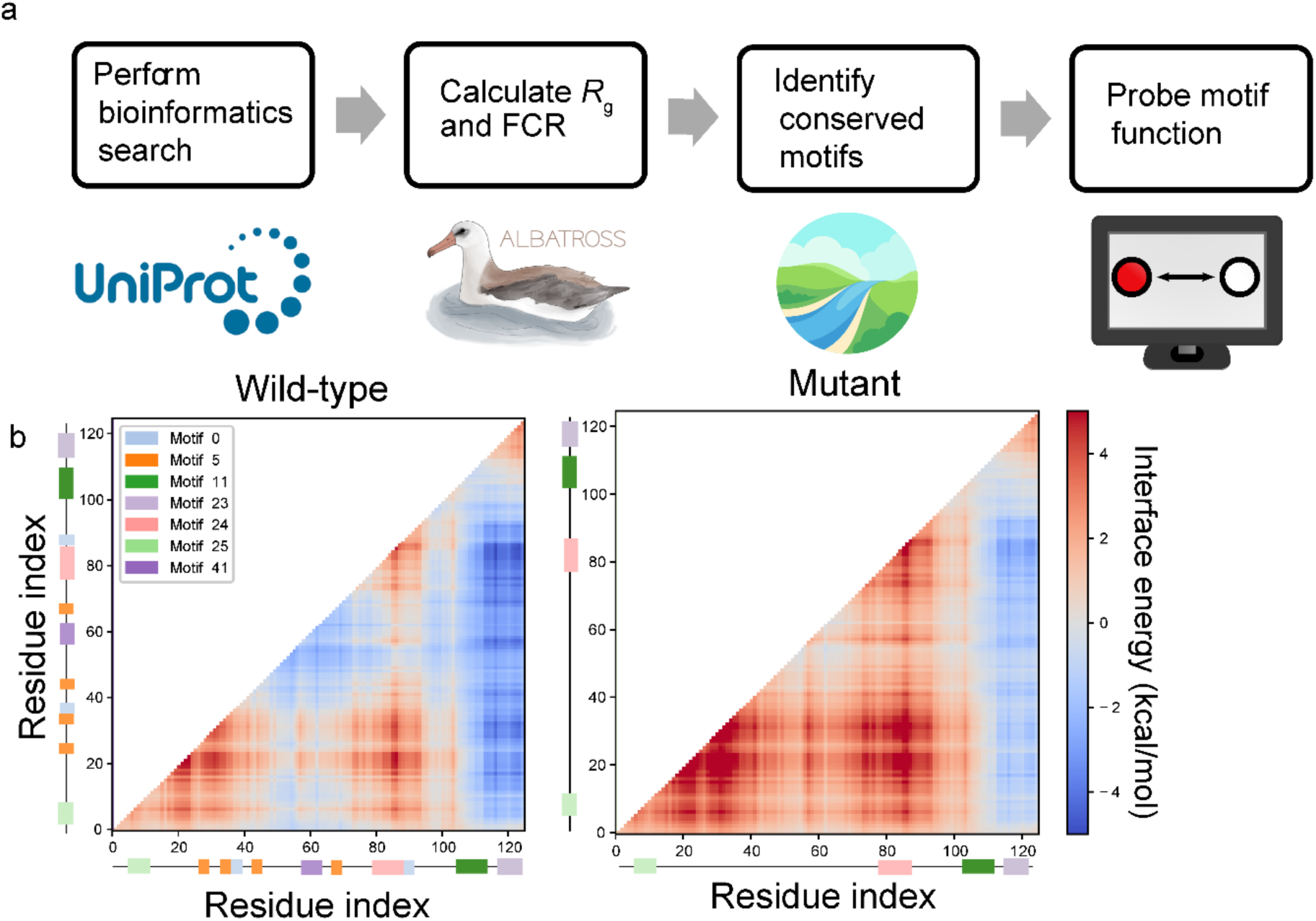
ML-driven algorithm identifies key sequence features. **(a)** Overview of computational methodology to identify repeated sequences in IDRs (motifs) that may be implicated in liquid-like phase separation behavior or ability to recruit small molecules. **(b)** Classified Dbp1n motifs were found to drive favorable protein-protein interface energies via FINCHES. A mutated Dbp1n sequence where motifs were disrupted showed unfavorable energies of interaction between homopolymeric protein chains.

**Extended Data Figure 2.**
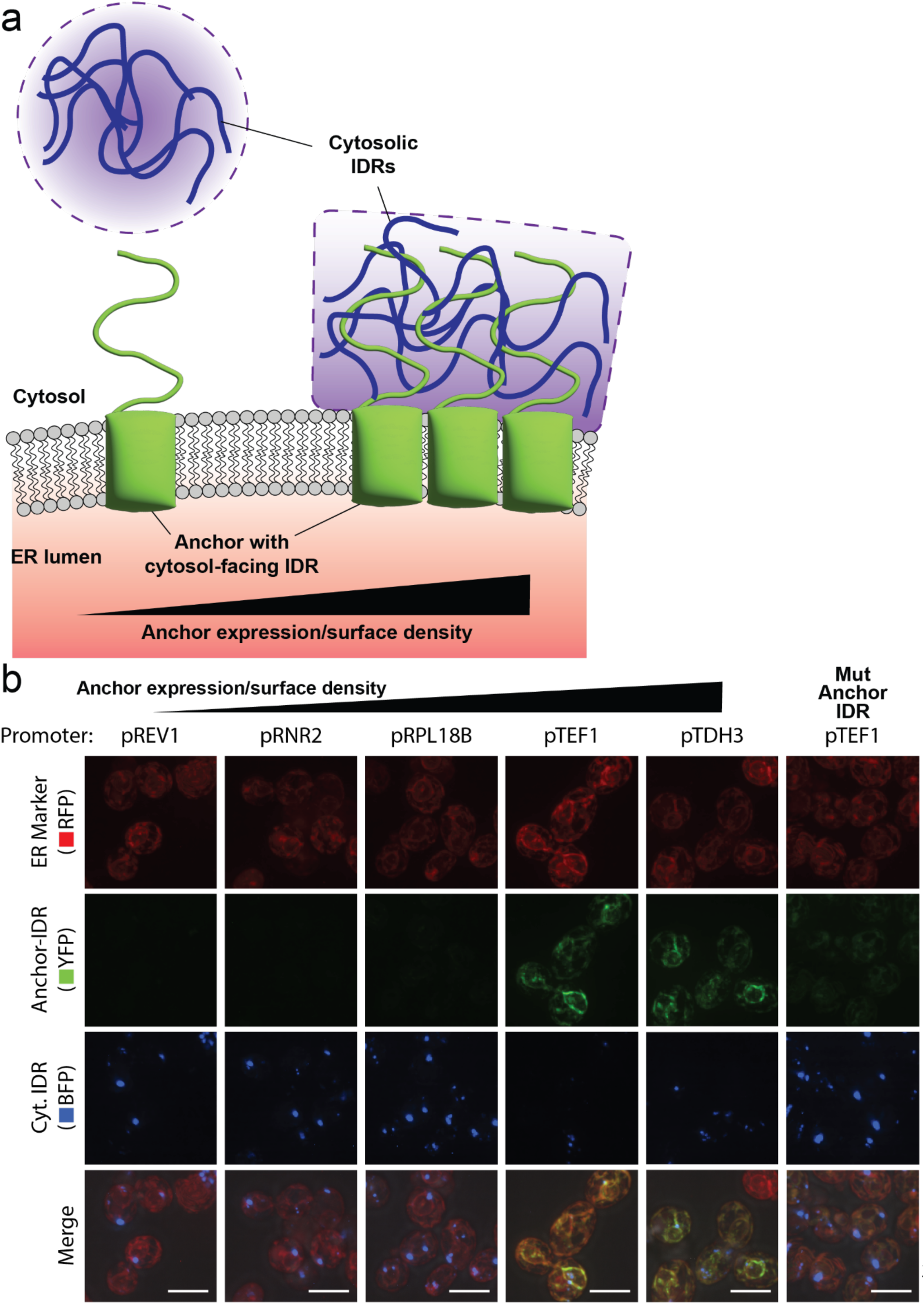
Formation of sMACs on the ER. **(a)** Co-expression of fluorescently labeled membrane-anchored and cytosolic IDRs results in merging of the cytosolic droplet onto the anchored structure. This ability to fuse with cytosolic droplet is used to demonstrate the liquid-like behaviors of sMACs. **(b)** Confocal microscopy of strains expressing increasing levels of Dbp1n-Ubc6p(233-250) fusion (left to right, first five columns, labeled with Venus YFP 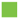) driven by constitutive promoters, along with a cytosolic trimeric coiled-coil Dbp1n (labeled with mTagBFP2 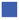) fusion. The rightmost column displays the negative control strain expressing Mut Dbp1n-Ubc6p(233-250) under pTEF1 control. Note the emergence of the YFP-labeled structures on the ER (labeled with mCherry RFP 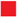) with pTEF1 and pTDH3 promoters with concomitant decrease in visible BFP-labeled cytosolic droplets and merging of small blue puncta onto the ER. This decrease in cytosolic BFP droplets was reversed when Mut Dbp1n was used to prevent phase separation and formation of sMACs in the membrane-anchored fusion (rightmost column), suggesting that cytosolic Dbp1n droplets were no longer merging onto the ER membrane. All scale bars represent 5 µm.

**Extended Data Figure 3.**
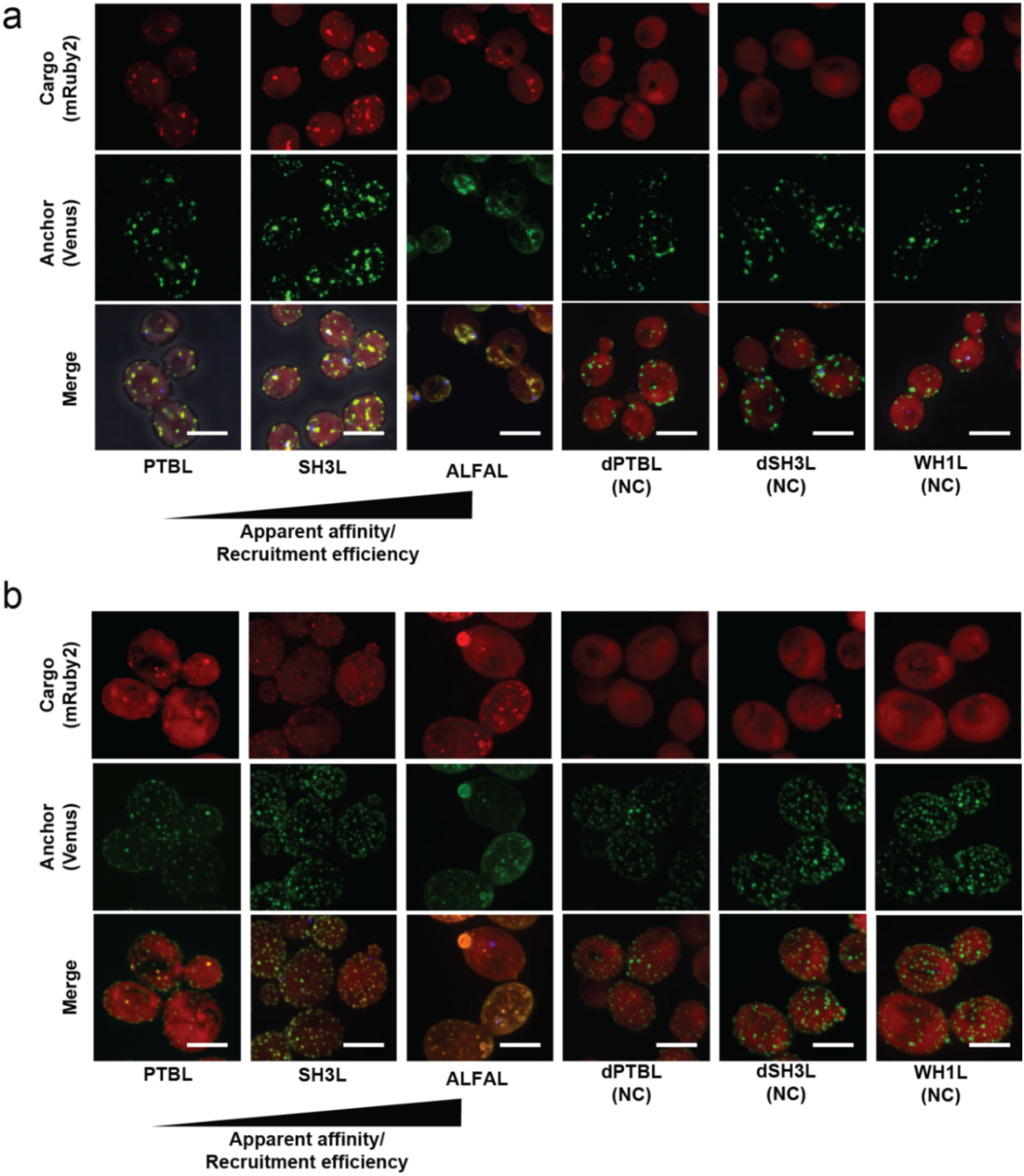
PpID-dependent recruitment of cargo into sMACs. **(a)** Recruitment of mRuby2 cargo into ER sMACs using different peptide ligands. **(b)** Recruitment of mRuby2 cargo into PM sMACs using different peptide ligands. As the apparent affinity of the PpID used for recruitment increases (see Extended Data Table 1), the efficiency of recruitment to sMACs also increase, similar to previous study that characterized cytosolic condensates with PpIDs^25^. Unexpectedly, however, the recruitment of ALFA ligand tagged cargo appeared to affect the morphology and abundance of sMACs on either the ER or PM. This follows previous observations that ALFA nanobody, when bound to its ligand, may be unstable in cells^107,108^. “NC” denotes negative controls using non-binding peptides. All scale bars represent 5 µm.

**Extended Data Figure 4.**
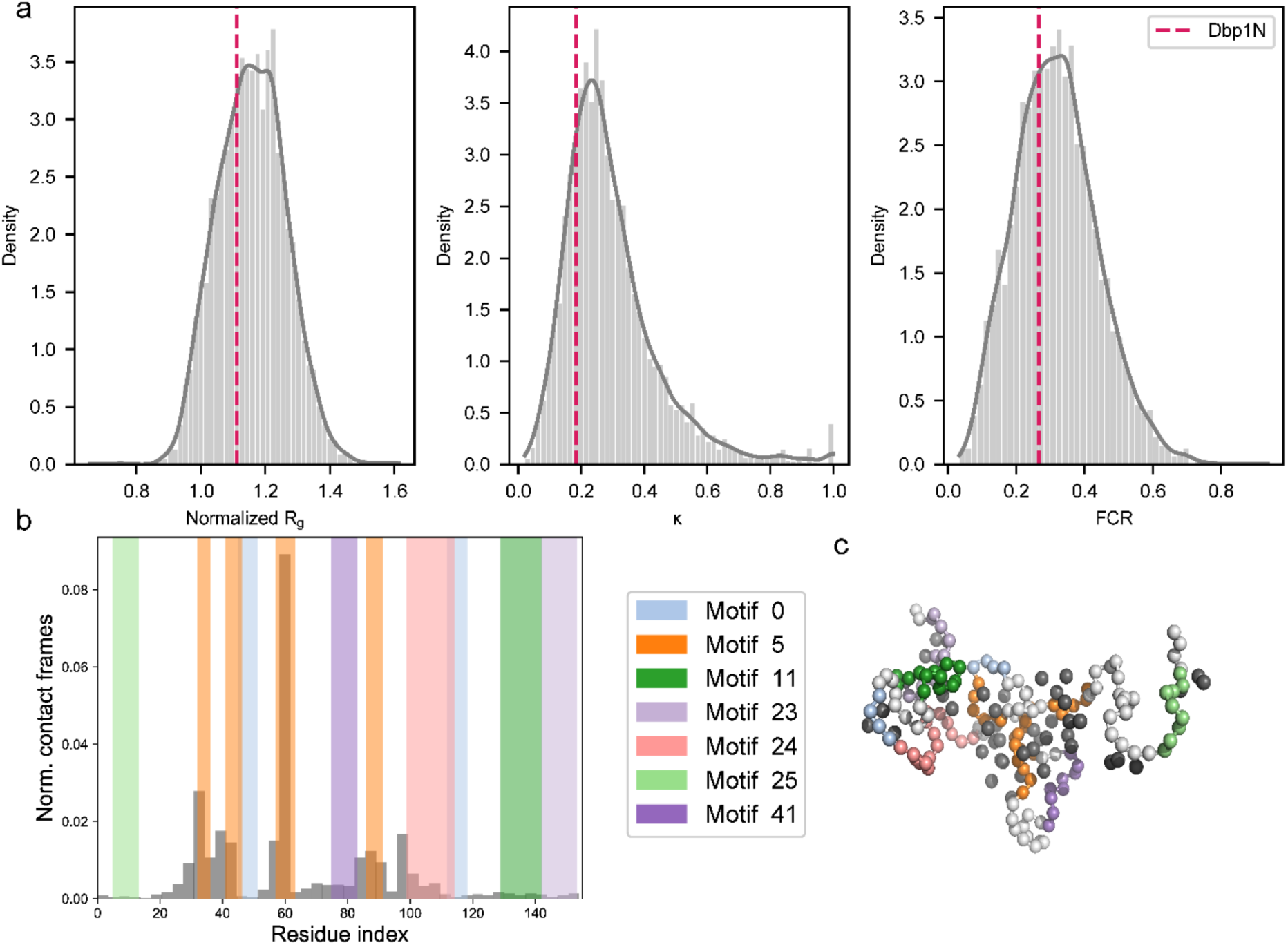
Dbp1n possesses several key characteristics of ATP-dependent RNA helicases that may be implicated in NAD(P)H recruitment. **(a)** Dbp1n is representative of the larger class of ATP-dependent RNA helicases in several single-chain properties associated with phase separation and potential NADH recruitment as calculated with the deep learning method ALBATROSS^74^. Together, these properties (radius of gyration (R_g_) normalized by the radius of gyration of an Analytical Flory Random Coil of the same number of amino acids, charge blockiness (κ), and fractional charge per residue (FCR)) point to characteristics of IDRs that may have affinity for NAD(P)H, which contains an adenosine moiety, phosphate(s), and a nicotinamide ring. (**b)** Enriched motifs extracted from ATP-dependent RNA helicases present in Dbp1n drive contacts between protein and NADH in single protein chain chain coarse-grained molecular dynamics simulation. Histogram is normalized such that the total area is equal to 1. (**c)** Representative pose of Dbp1n—NADH contacts from single-chain simulation. The Dbp1n chain is shown as white spheres, and motifs found along the Dbp1n sequence are colored according to the legend to the left. NADH is represented as three gray spheres.

**Extended Data Figure 5.**
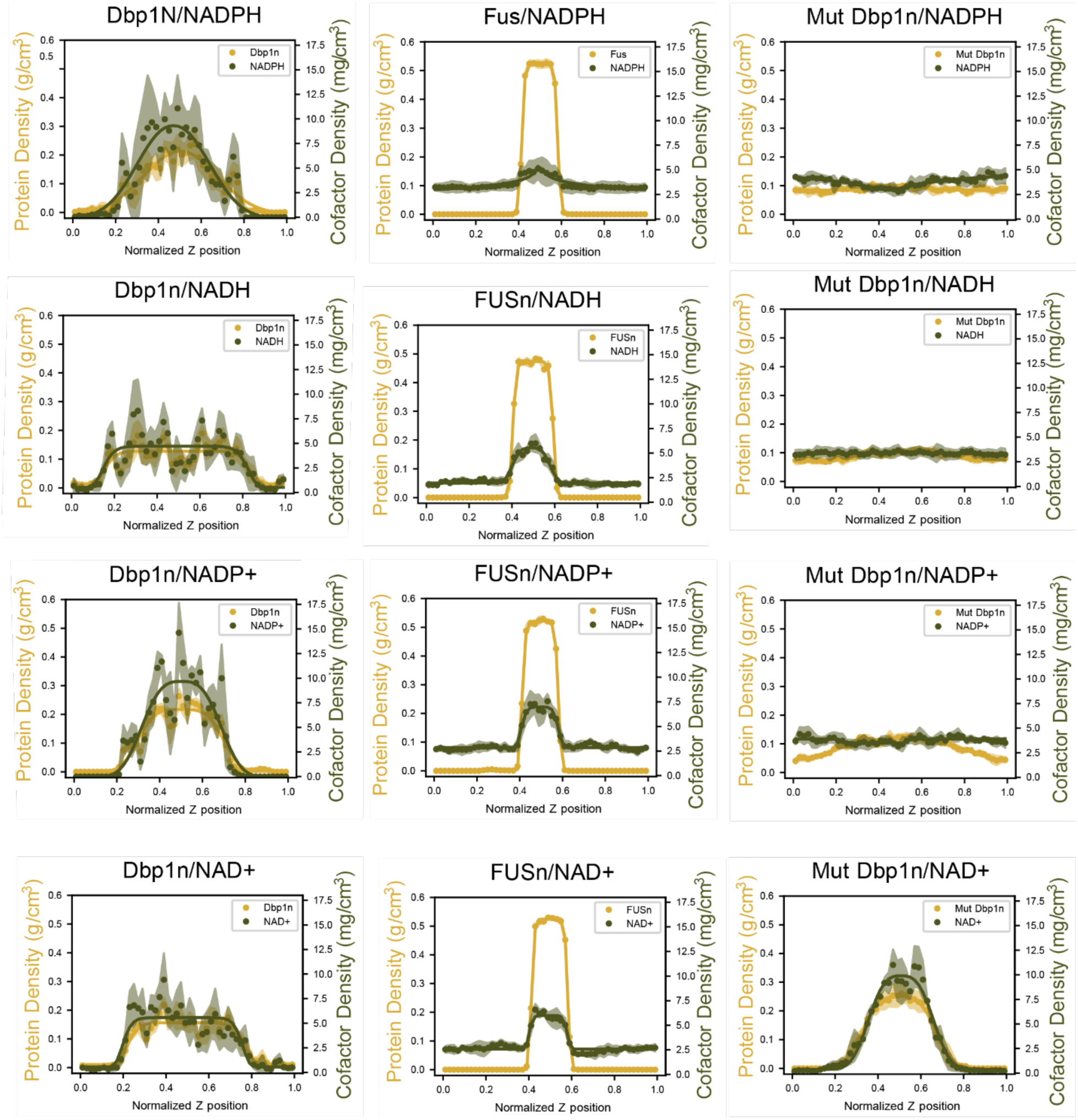
Molecular dynamics simulations show differential partitioning of cofactors into Dbp1n but not FUSn. Points represent the time-averaged density over equilibrated simulation frames at a given Z-axis position, shaded region represents the standard error, and curve represents a super-Gaussian fit to data.

**Extended Data Figure 6.**
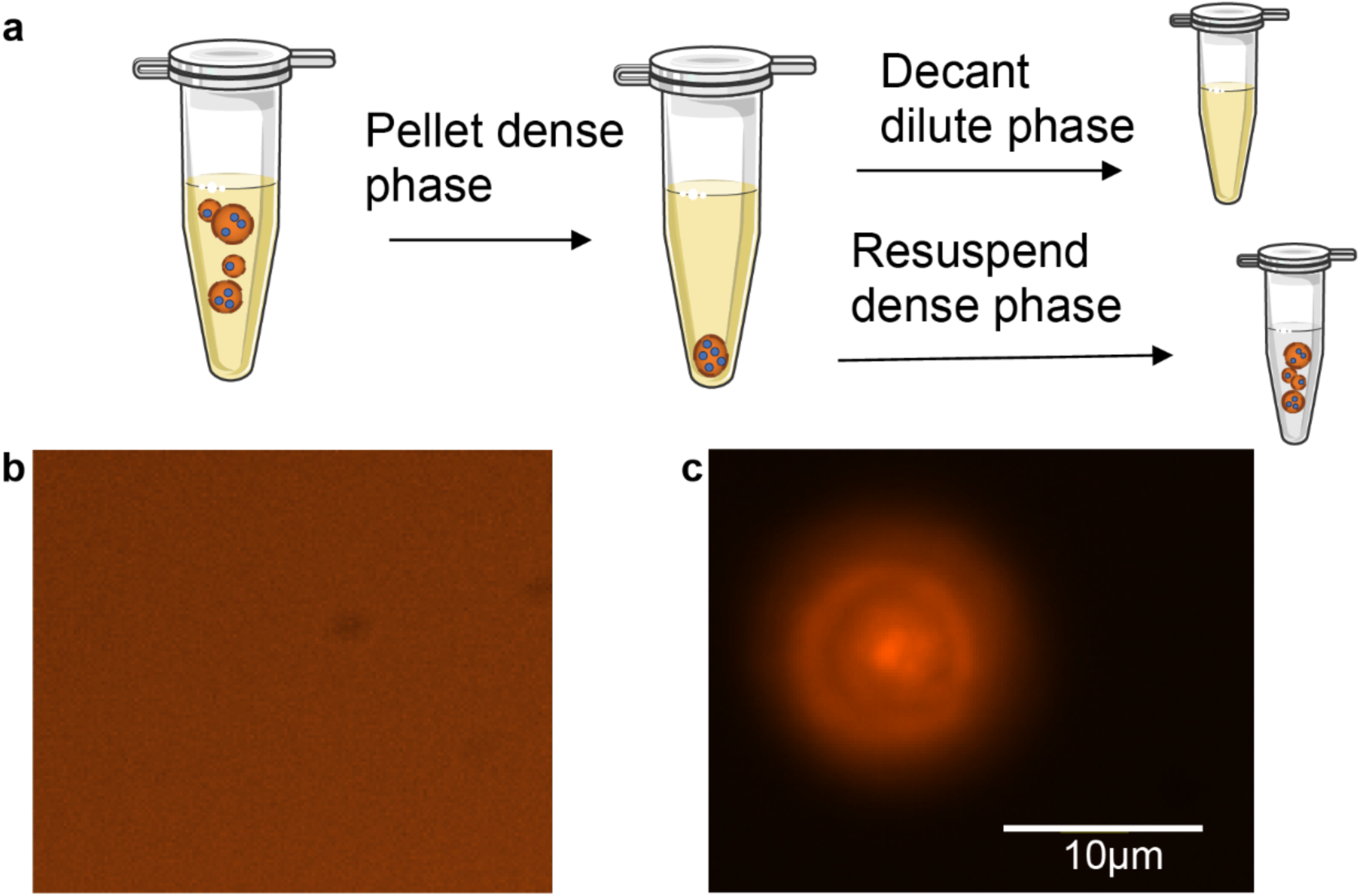
Experimental validation of NADPH and NADH partitioning into purified Dbp1n condensates *in vitro*. **(a)** Solutions of purified Dbp1n IDR fused to mCherry were combined with either NADH or NADPH. The dense phase was pelleted via centrifugation and mechanically separated from the dilute phase, then NADH and protein concentration was quantified via fluorescence. **(b)** Widefield microscopy of Dbp1n fused to mCherry and trimeric coiled-coil domain in supercritical solution conditions (3M urea, 1M NaCl), imaged through fluorescence at 555 nm excitation. **(c)** Dbp1n fused to mCherry and trimeric coiled-coil domain formed droplets in subcritical solution conditions in widefield microscopy. Measured partition coefficients are tabulated in Table 1.

**Extended Data Table 1.**
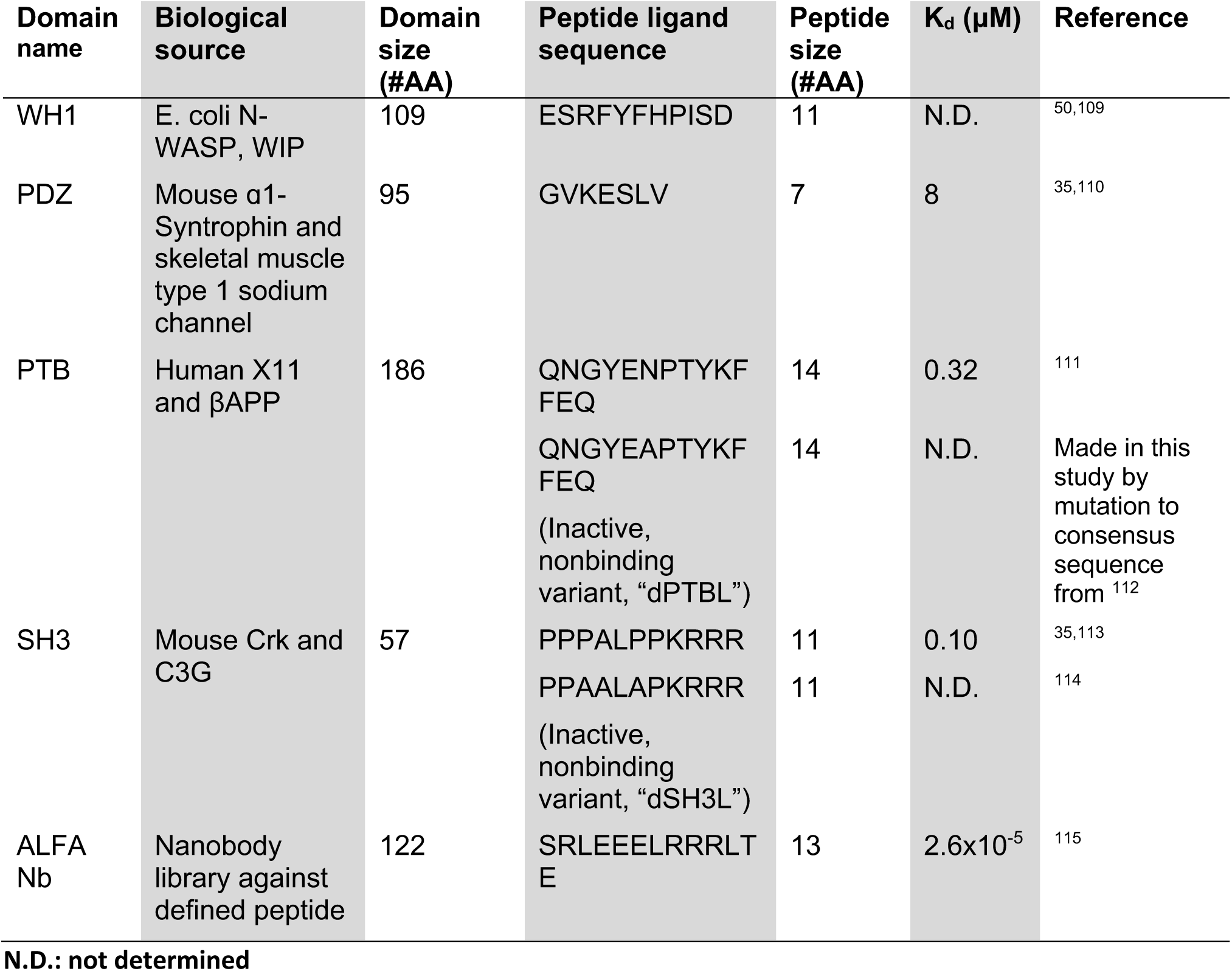
Characteristics of PpIDs used in the study, listed in order of decreasing reported Kd.

| Domain name | Biological source | Domain size (#AA) | Peptide ligand sequence | Peptide size (#AA) | $K_d$ (μM) | Reference |
| --- | --- | --- | --- | --- | --- | --- |
| WH1 | E. coli N-WASP, WIP | 109 | ESRFYFHPISD | 11 | N.D. | 50,109 |
| PDZ | Mouse α1-Syntrophin and skeletal muscle type 1 sodium channel | 95 | GVKESLV | 7 | 8 | 35,110 |
| PTB | Human X11 and βAPP | 186 | QNGYENPTYKF<br>FEQ | 14 | 0.32 | 111 |
|  |  |  | QNGYEAPTYKF<br>FEQ<br>(Inactive, nonbinding variant, "dPTBL") | 14 | N.D. | Made in this study by mutation to consensus sequence from <sup>112</sup> |
| SH3 | Mouse Crk and C3G | 57 | PPPALPPKRRR | 11 | 0.10 | 35,113 |
|  |  |  | PPAALAPKRRR<br>(Inactive, nonbinding variant, "dSH3L") | 11 | N.D. | 114 |
| ALFA Nb | Nanobody library against defined peptide | 122 | SRLEELRRRLT<br>E | 13 | $2.6 \times 10^{-5}$ | 115 |

**Extended Data Table 2.**
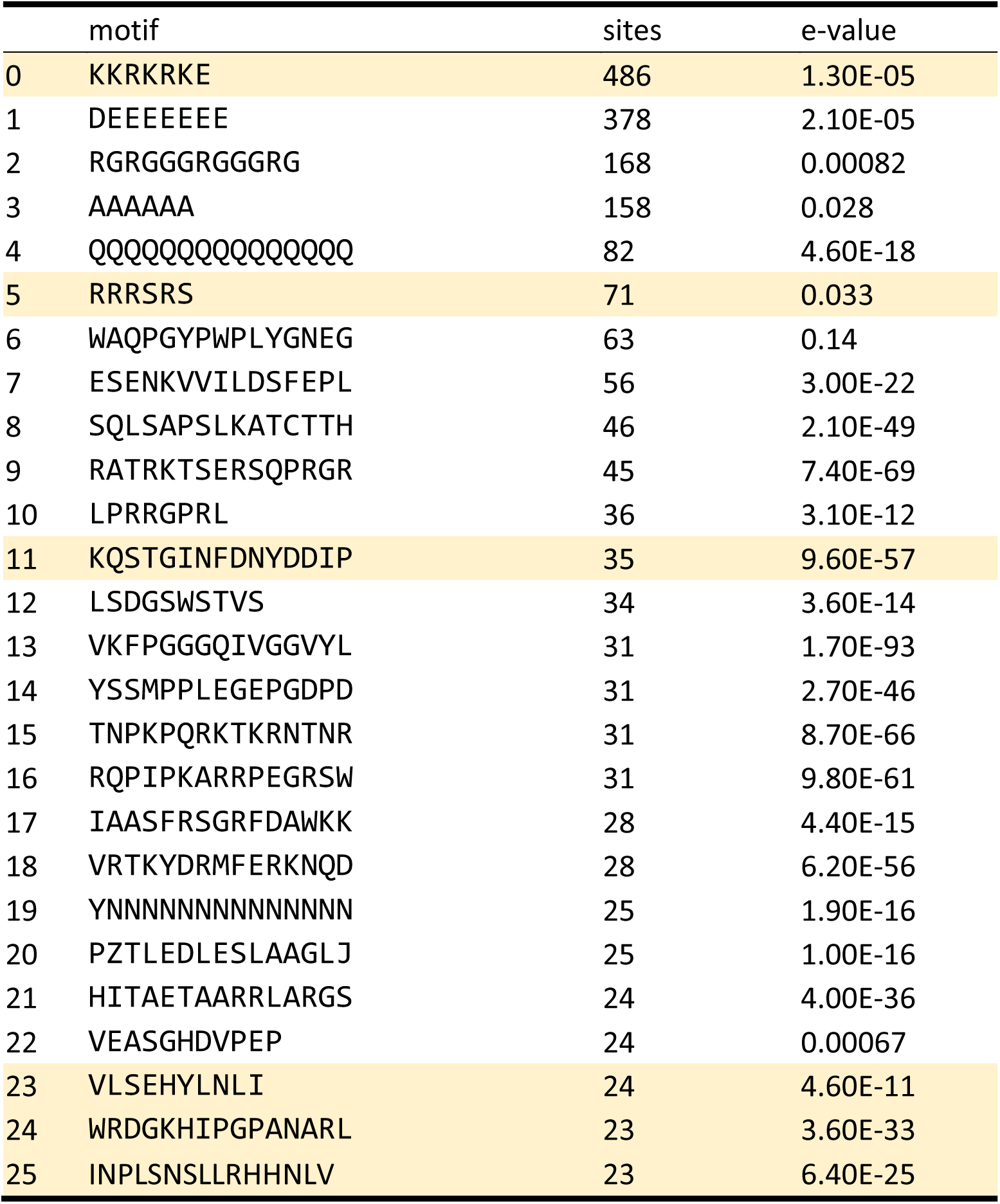
List of top enriched IDR motifs identified by sequence screen with affinity for the adenosine moiety or phosphate group in ATP-dependent RNA helicases. Sites represent the number of times a motif appears in the bioinformatic database, and e-value is a measure of probability of that sequence occurring due to random chance. Highlighted motifs (with point mutations tolerated using a BLOSUM sequence similarity tolerance of 75%) in Dbp1n. None of these motifs appear in FUSn.

## Supplementary Information

### Supplementary Note 1. Compositions and amino acid sequences of sMAC building blocks

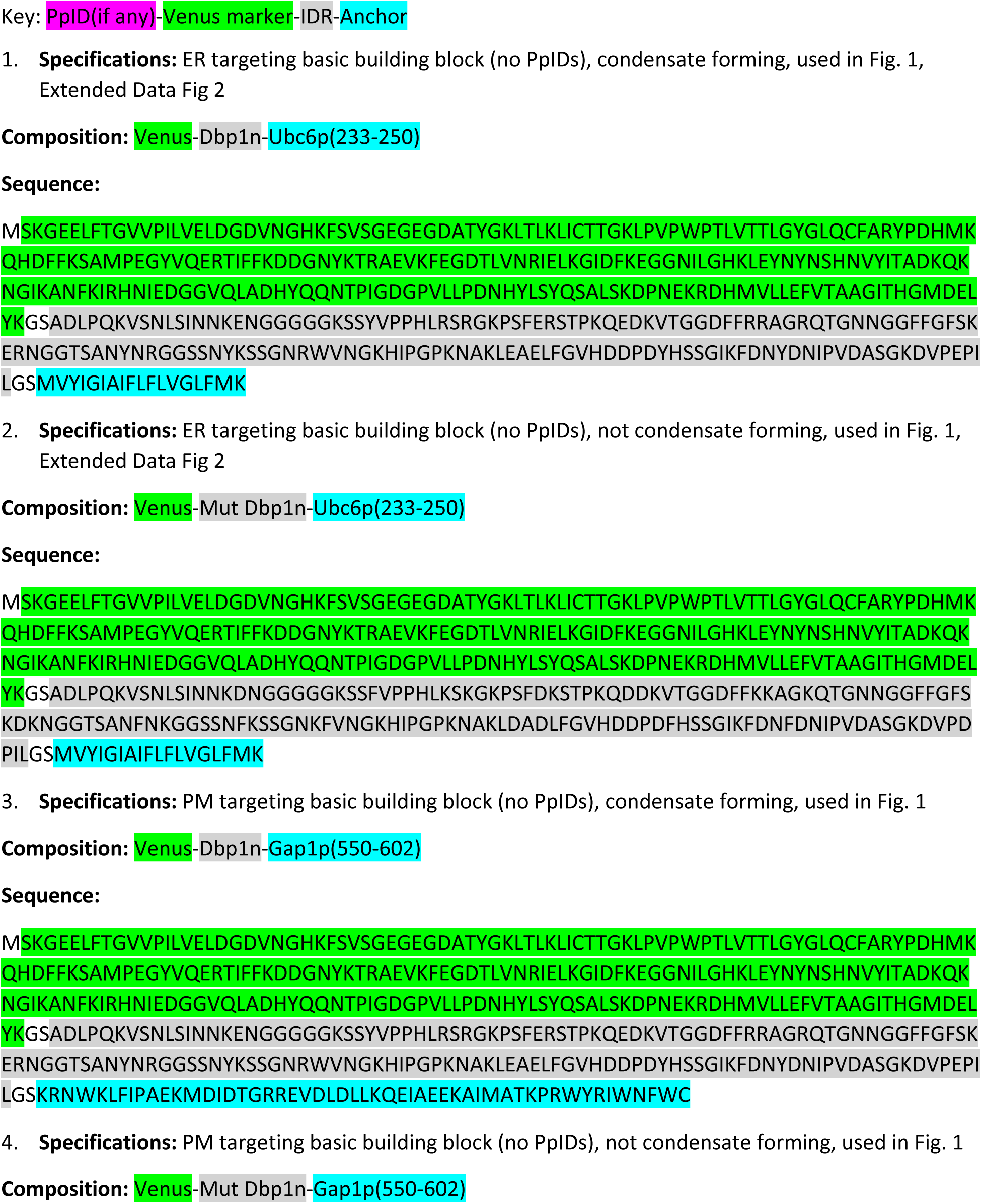
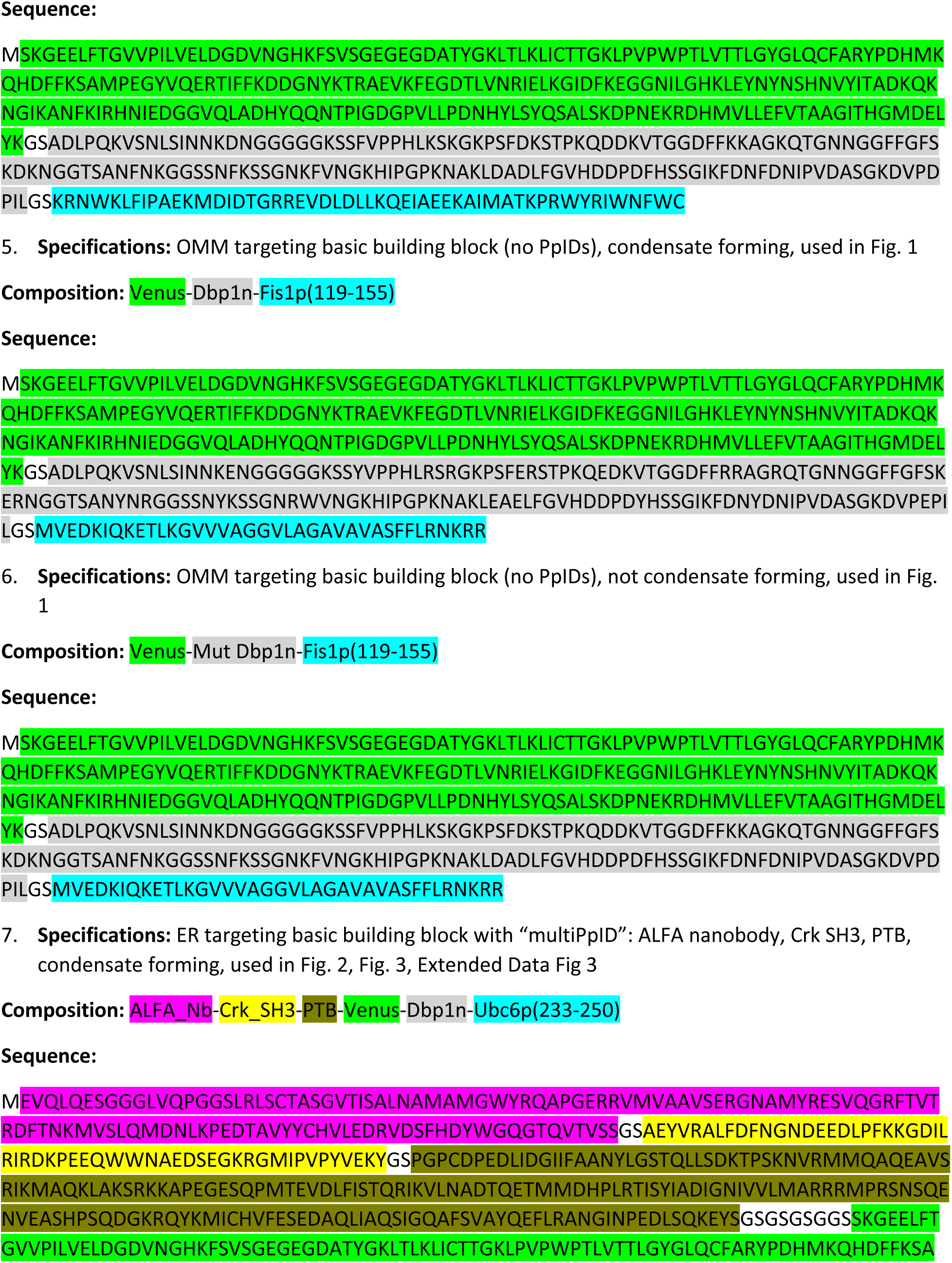
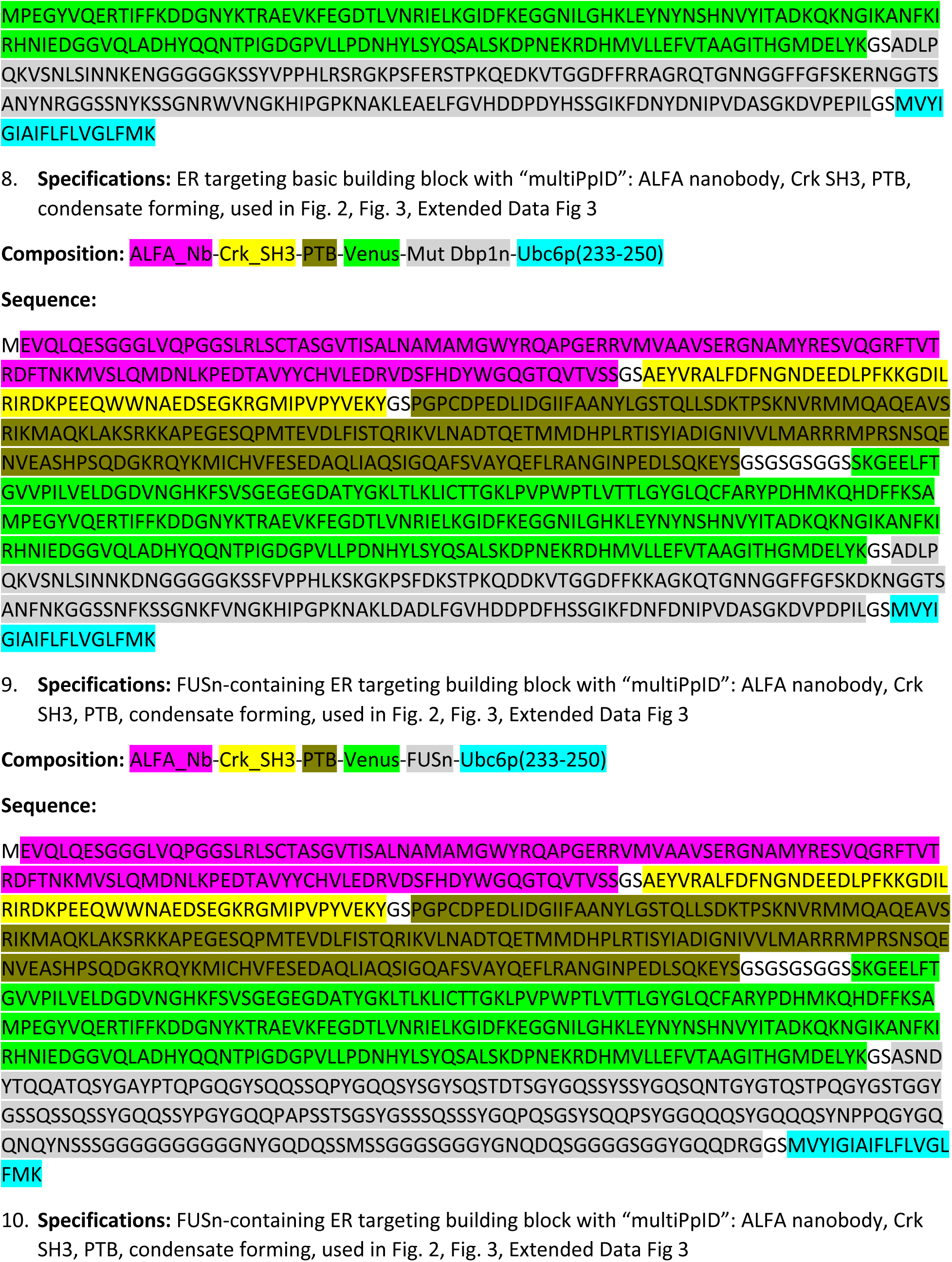
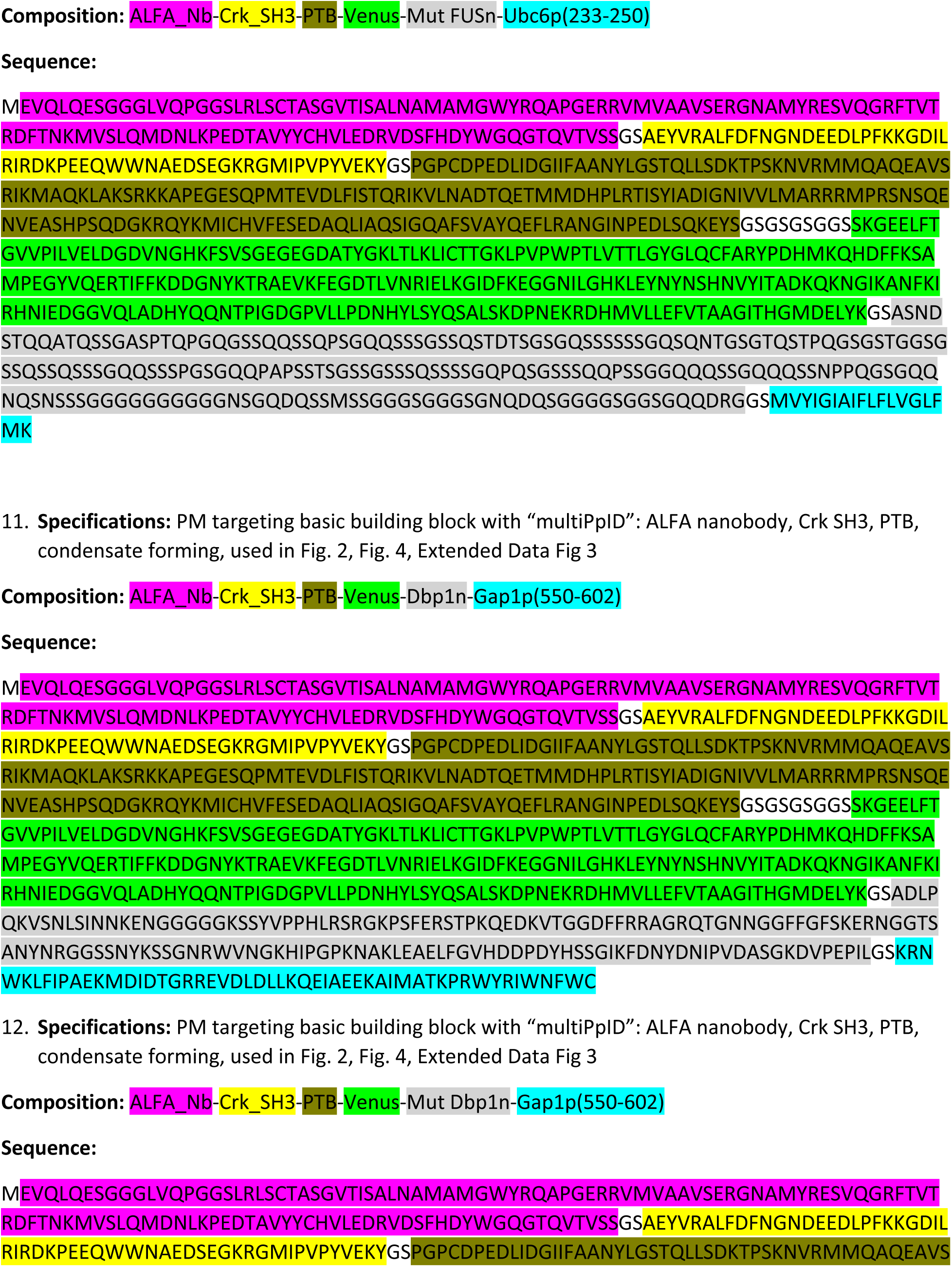
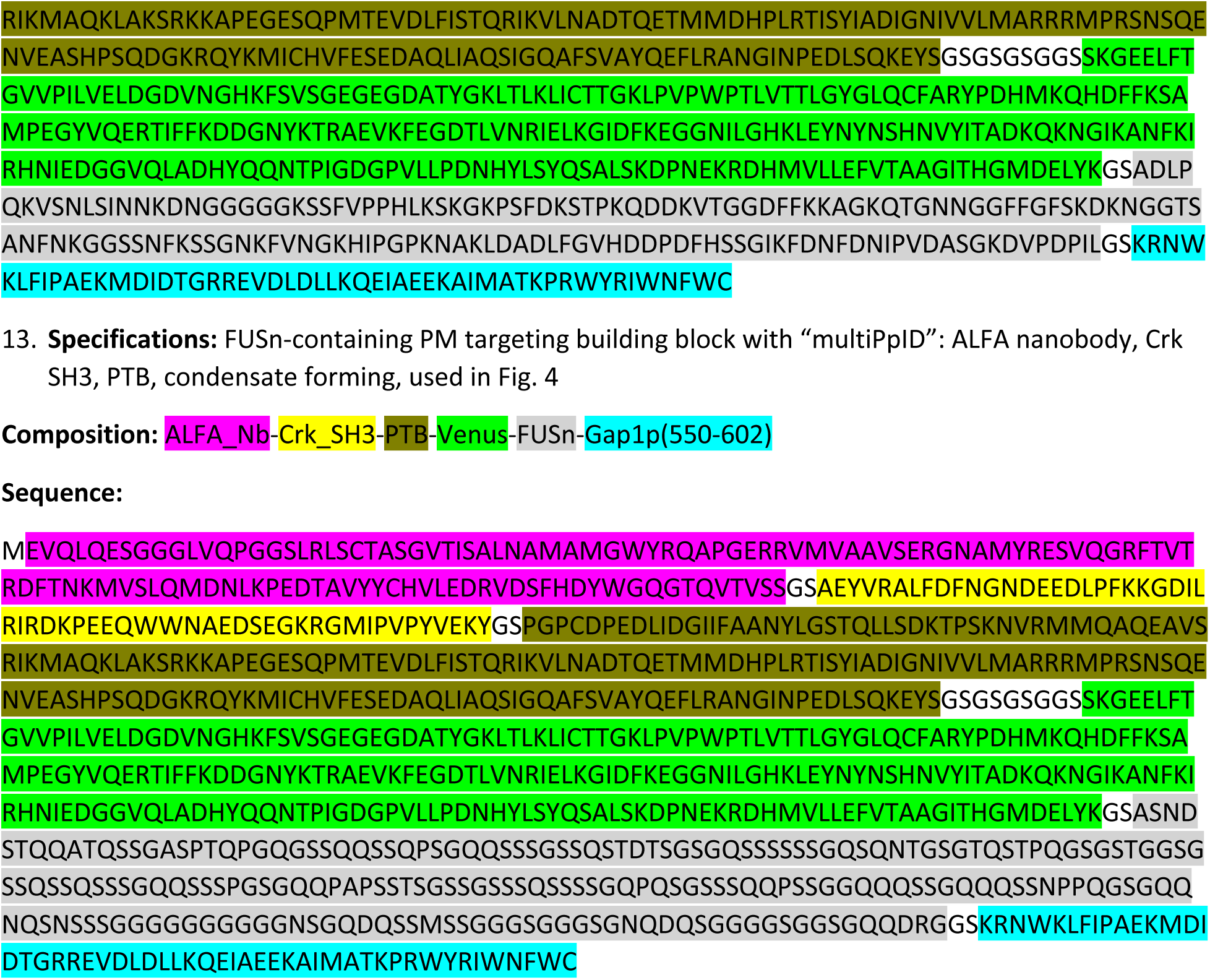

### Supplementary Note 2. Formation of cytosolic condensates by Dbp1n using engineered oligomerization domains

In contrast to the defined, high-affinity interfaces characteristic of folded protein complexes, intrinsically disordered regions (IDRs) such as Dbp1n assemble through a multitude of weak, transient interactions distributed along the polypeptide chain. The cumulative effect of these low-affinity interactions, referred to as multivalency, is a primary determinant of the thermodynamic propensity for phase separation. Repetition of short sequence motifs within an IDR increases the number of potential contact sites, thereby strengthening both homotypic and heterotypic interactions and lowering the free-energy barrier to condensate formation. Multivalency encoded at the sequence level directly modulates the saturation concentration for condensation and governs key material properties of the dense phase, including its viscosity, elasticity, and coarsening dynamics.

To further promote condensation under physiological conditions, multivalency can be enhanced synthetically by genetically encoding tandem repeats of IDR sequences or introducing oligomerization domains. In this study, cytosolic condensates were engineered to include a homo-trimeric coiled-coil domain^116^, which increases the effective valency of Dbp1n and promotes robust phase separation in *S. cerevisiae* (Supplementary Fig 1).

**Supplementary Figure 1.**
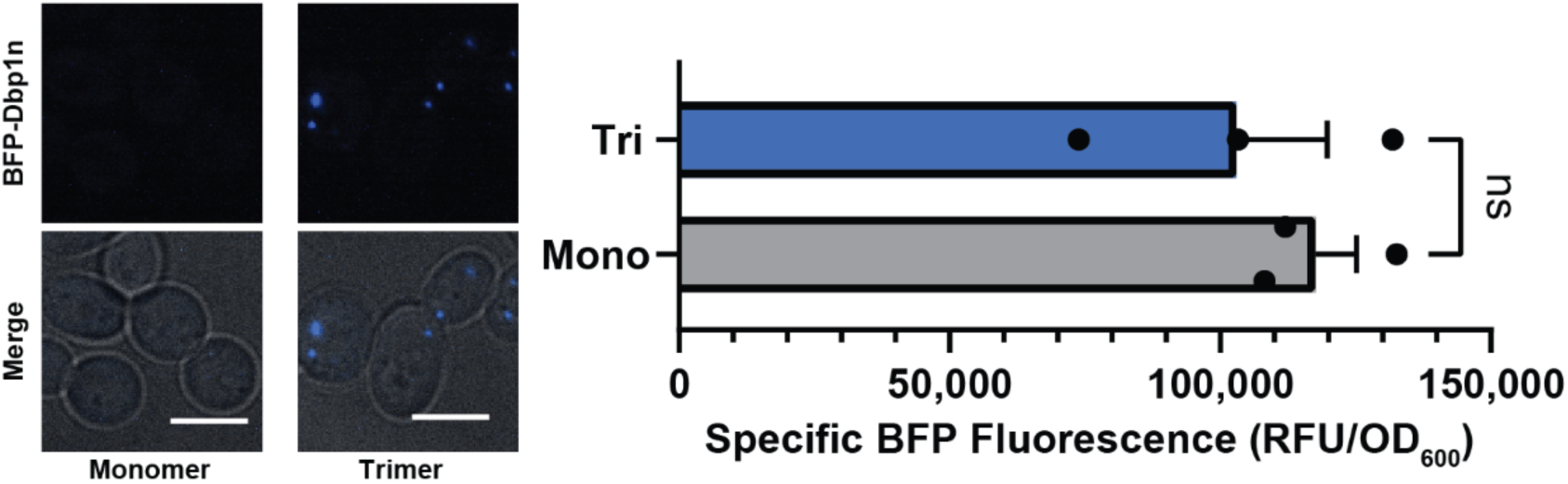
Cytosolic Dbp1n condensates only forms when IDR is fused to an oligomerization domain. Under the control of identical expression cassette with a pCCW12 promoter on a low-copy number plasmid, BFP-labeled Dbp1n formed visible cytosolic puncta only when fused to a homo-trimeric coiled coil domain (5L6HC3_1^116^) that enhanced multivalency but did not in the absence of an oligomerization domain. “Tri” denotes trimeric Dbp1n; “Mono” denotes monomeric Dbp1n. All scale bars represent 5 µm. Two-tailed Student’s *t*-test was used to compare the sample data between each group of three biological replicates. All data are shown as the mean ± s.e.m. ns denotes not significant.

**Supplementary Figure 2.**
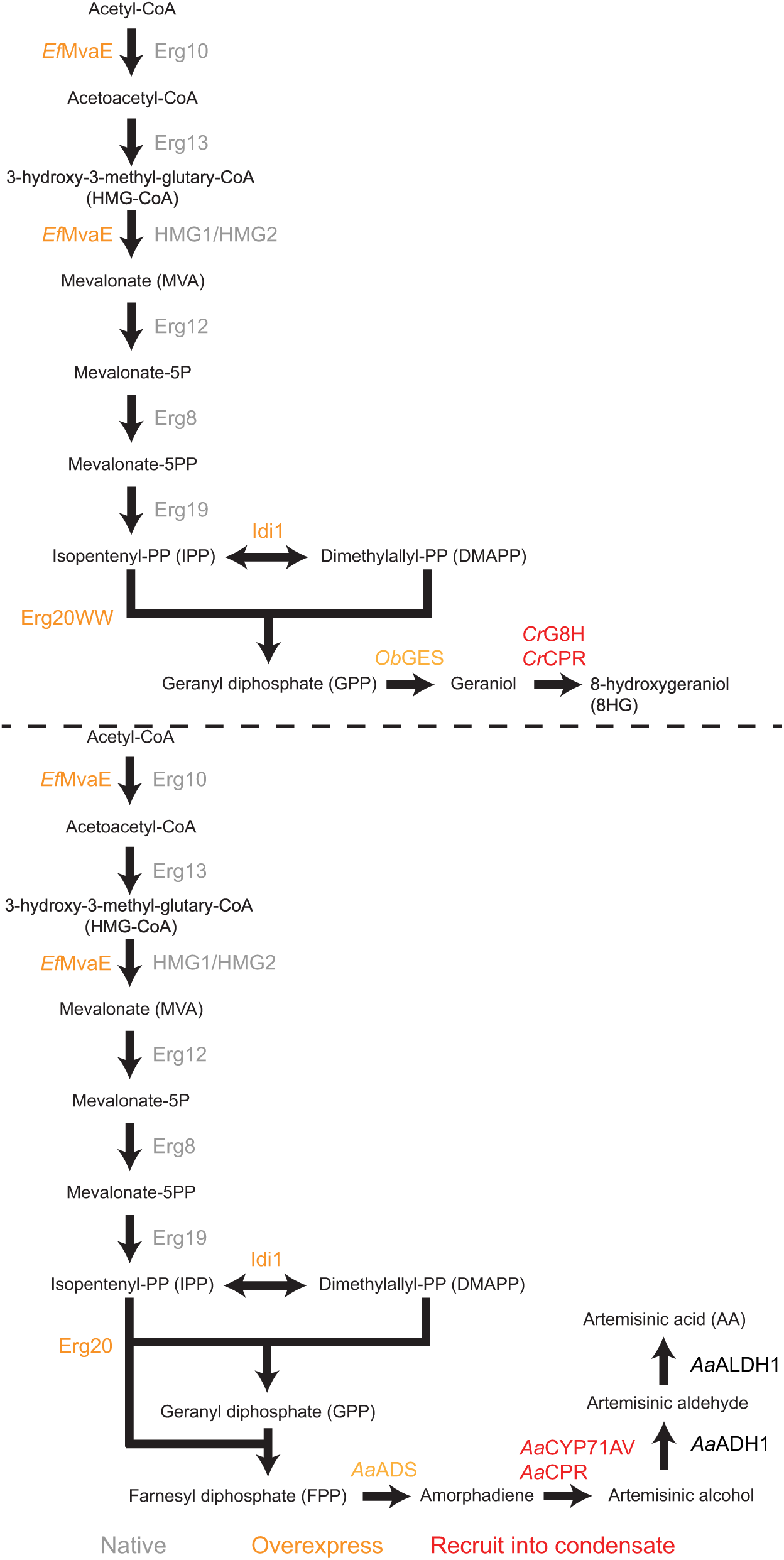
8-hydroxygeraniol (8HG) and artemisinic acid (AA) biosynthetic pathways introduced into *S. cerevisiae* strains used in the study. Biosynthesis of both 8HG and AA share the diphosphate precursors (IPP and DMAPP) from the mevalonate (MVA) pathway. To boost the level of the precursors, known rate-limiting enzymatic steps of the native MVA pathway were supplemented by the overexpression of two corresponding enzymes (*Ef*MvaE and Idi1). The specific terpenoid scaffolds for 8HG and AA (geraniol and amorphadiene, respectively) were synthesized by the overexpressed GPP-specific Erg20WW mutant and Erg20, followed by their respective synthases *Ob*GES and *Aa*ADS. Both terpenoid scaffolds were then subjected to cytochrome P450-driven catalysis for hydroxylation, the efficiency of which might be improved by recruitment into sMACs.

**Supplementary Table 1.**
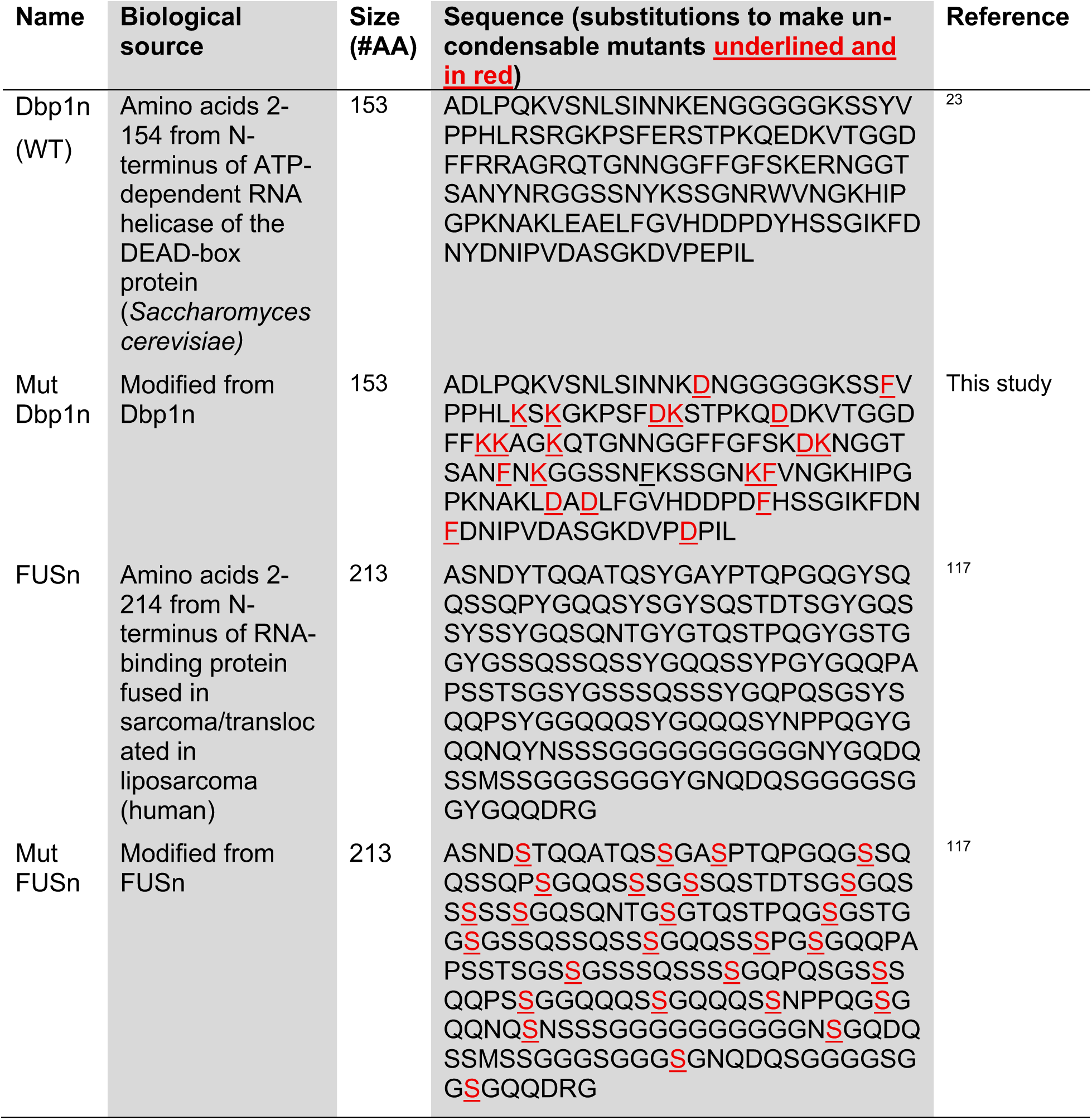
IDR amino acid sequences used in the study.

**Supplementary Table 2.**
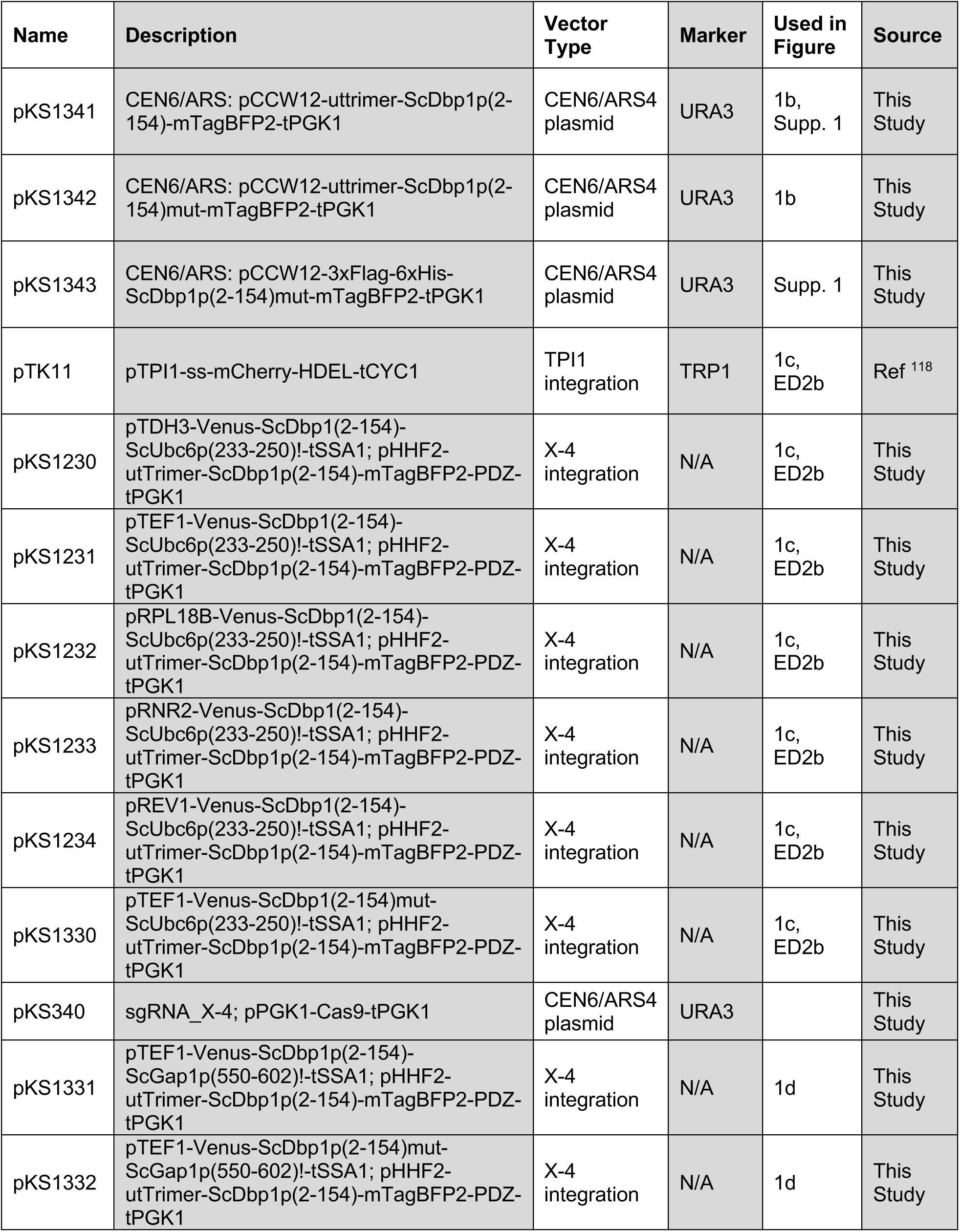
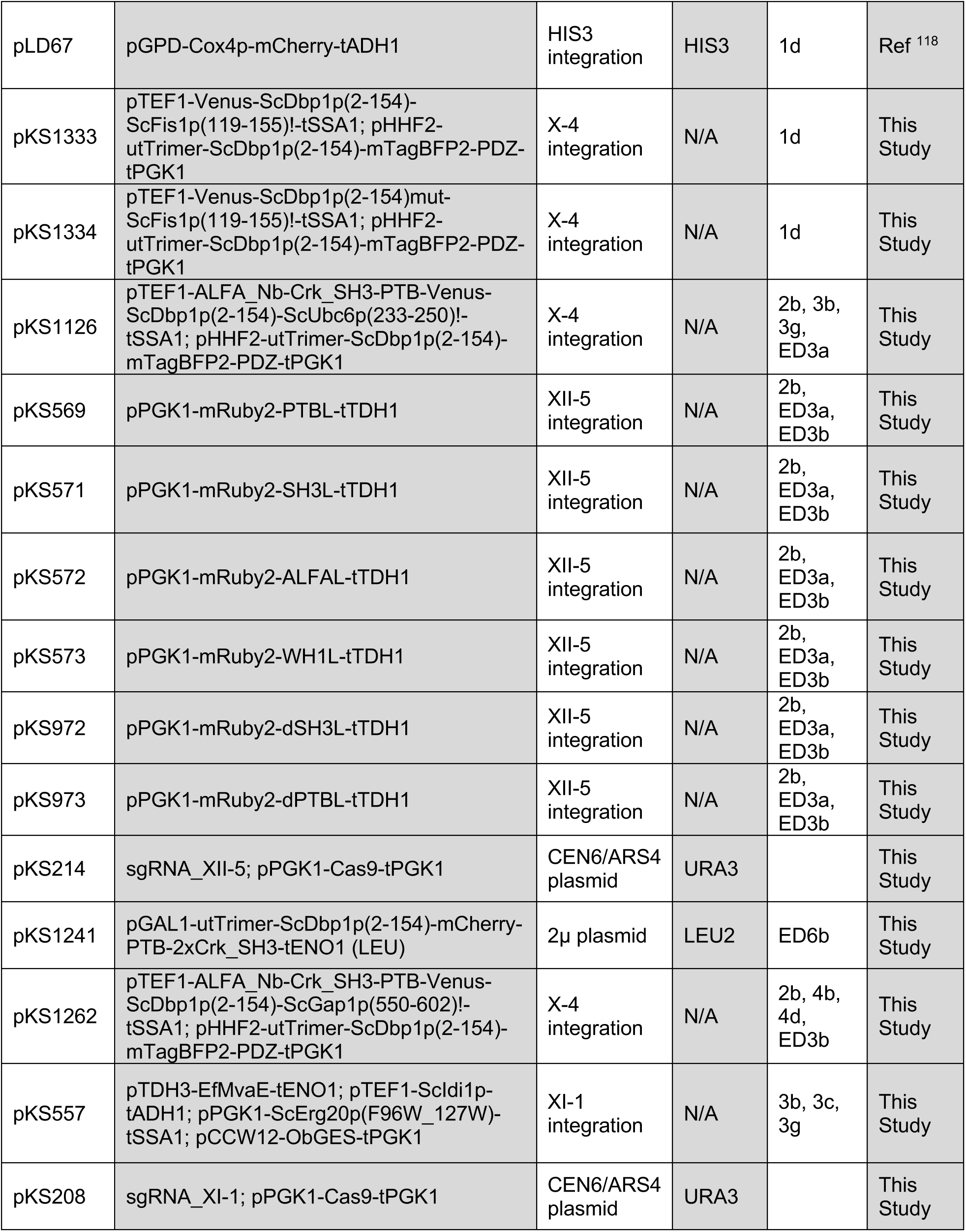
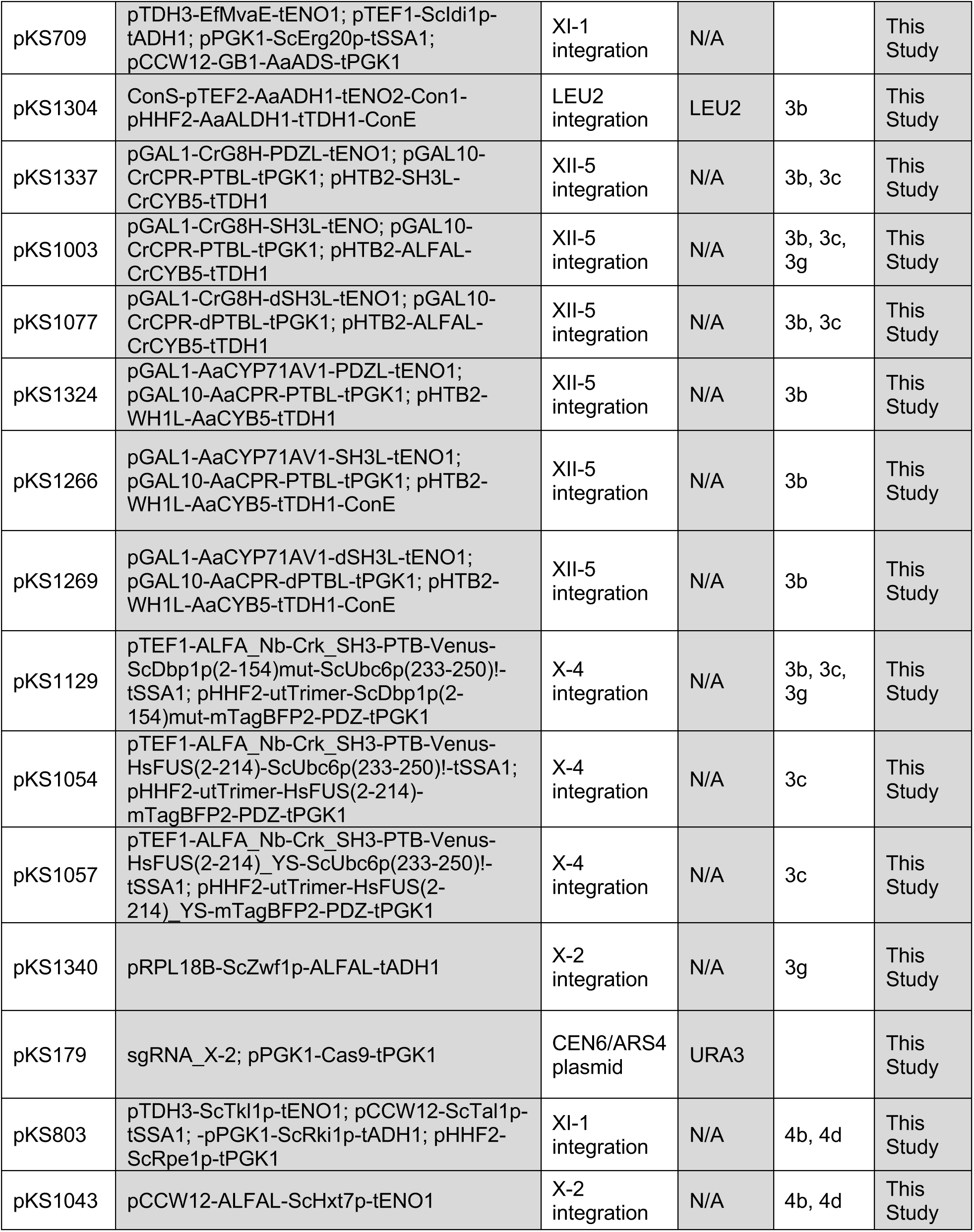
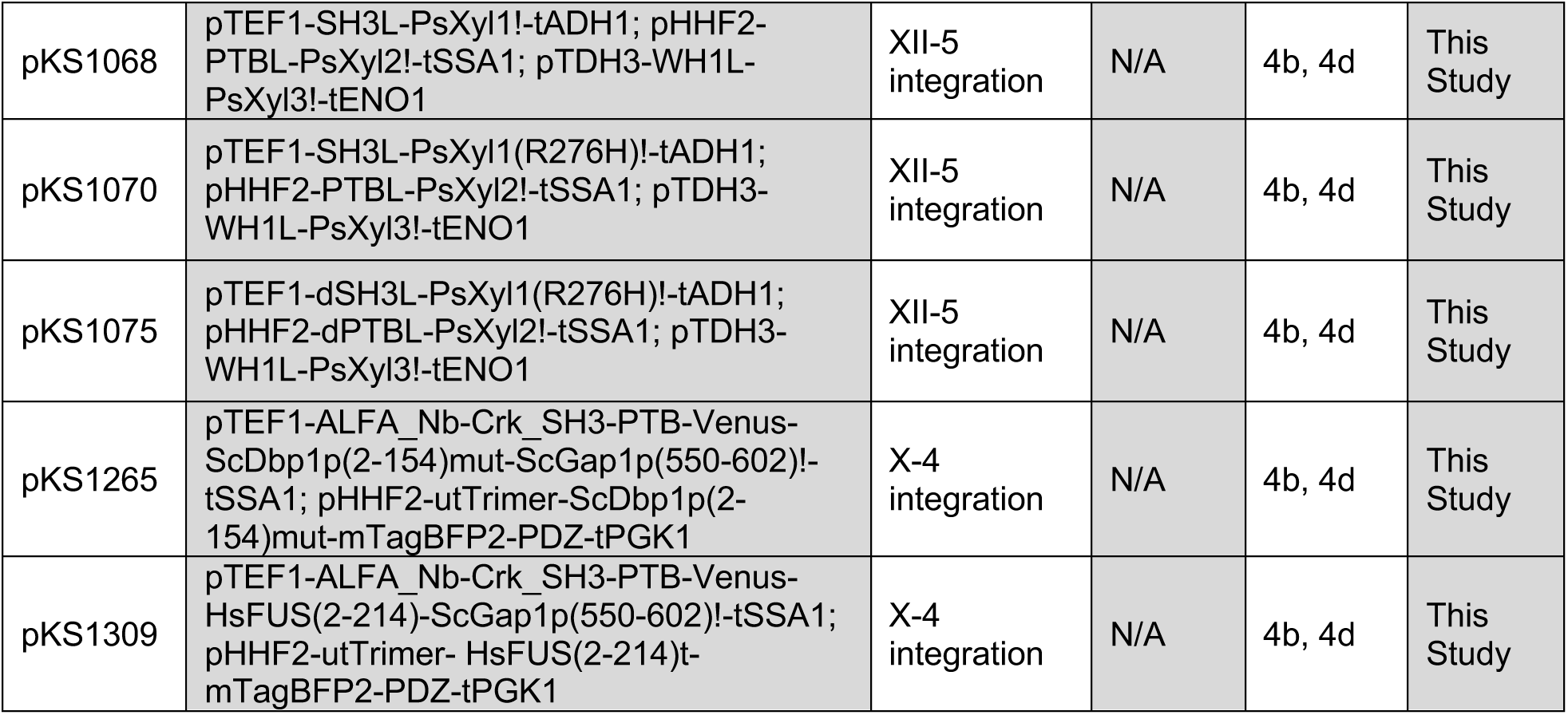
Plasmids used in study.

**Supplementary Table 3.** Yeast strains used in the study.

| Name | Genotype | Parent | Plasmids used for construct | Marker(s) | Used in Figure | Source |
| --- | --- | --- | --- | --- | --- | --- |
| CEN.PK2-1C | MATa ura3-52 trp1-289 leu2-3,112 his3Δ1 MAL2-8C SUC2 | N/A, common lab strain |  | N/A |  | Avalos Lab |
| yKS418 | CEN.PK2-1C CEN6/ARS: pCCW12-uttrimer-ScDbp1p(2-154)-mTagBFP2-tPGK1 | CEN.PK2-1C | pKS1341 | URA3 | 1b, Supp. 1 | This Study |
| yKS419 | CEN.PK2-1C CEN6/ARS: pCCW12-uttrimer-ScDbp1p(2-154)mut-mTagBFP2-tPGK1 | CEN.PK2-1C | pKS1342 | URA3 | 1b | This Study |
| yKS420 | CEN.PK2-1C CEN6/ARS: pCCW12-3xFlag-6xHis-ScDbp1p(2-154)-mTagBFP2-tPGK1 | CEN.PK2-1C | pKS1343 | URA3 | Supp. 1 | This Study |
| yKS019 | CEN.PK2-1C pTPI1::pTPI1-ss-mCherry-HDEL-tCYC1 | CEN.PK2-1C | pTK11 | TRP1 |  | This Study |
| yKS372 | yKS019 X-4:: pTDH3-Venus-ScDbp1(2-154)-ScUbc6p(233-250)!-tSSA1-Con1-pHHF2-utTrimer-ScDbp1p(2-154)-mTagBFP2-PDZ-tPGK1 | yKS019 | pKS1230 | TRP1 | 1c, ED2b | This Study |
| yKS373 | yKS019 X-4:: pTEF1-Venus-ScDbp1(2-154)-ScUbc6p(233-250)!-tSSA1-Con1-pHHF2-utTrimer-ScDbp1p(2-154)-mTagBFP2-PDZ-tPGK1 | yKS019 | pKS1231 | TRP1 | 1c, ED2b | This Study |
| yKS374 | yKS019 X-4:: pRPL18B-Venus-ScDbp1(2-154)-ScUbc6p(233-250)!-tSSA1-Con1-pHHF2-utTrimer-ScDbp1p(2-154)-mTagBFP2-PDZ-tPGK1 | yKS019 | pKS1232 | TRP1 | 1c, ED2b | This Study |
| yKS375 | yKS019 X-4:: pRNR2-Venus-ScDbp1(2-154)-ScUbc6p(233-250)!-tSSA1-Con1-pHHF2-utTrimer-ScDbp1p(2-154)-mTagBFP2-PDZ-tPGK1 | yKS019 | pKS1233 | TRP1 | 1c, ED2b | This Study |
| yKS376 | yKS019 X-4:: pREV1-Venus-ScDbp1(2-154)-ScUbc6p(233-250)!-tSSA1-Con1-pHHF2-utTrimer-ScDbp1p(2-154)-mTagBFP2-PDZ-tPGK1 | yKS019 | pKS1234 | TRP1 | 1c, ED2b | This Study |
| yKS377 | yKS019 X-4:: pTEF1-Venus-ScDbp1(2-154)mut-ScUbc6p(233-250)!-tSSA1-Con1-pHHF2-utTrimer-ScDbp1p(2-154)-mTagBFP2-PDZ-tPGK1 | yKS019 | pKS1330 | TRP1 | 1c, ED2b | This Study |
| yKS378 | CEN.PK2-1C X-4:: pTEF1-Venus-ScDbp1p(2-154)-ScGap1p(550-602)!-tSSA1-Con1-pHHF2-utTrimer-ScDbp1p(2-154)-mTagBFP2-PDZ-tPGK1 | CEN.PK2-1C | pKS1331 | TRP1 | 1d | This Study |
| yKS379 | CEN.PK2-1C X-4:: pTEF1-Venus-ScDbp1p(2-154)mut-ScGap1p(550-602)!-tSSA1-Con1-pHHF2-utTrimer-ScDbp1p(2-154)-mTagBFP2-PDZ-tPGK1 | CEN.PK2-1C | pKS1332 | TRP1 | 1d | This Study |
| yKS101 | CEN.PK2-1C HIS3::pGPD-Cox4p-mCherry-tADH1 | CEN.PK2-1C | pLD76 | HIS3 |  | This Study |
| yKS402 | yKS101 X-4:: pTEF1-Venus-ScDbp1p(2-154)-ScFis1p(119-155)!-tSSA1-Con1-pHHF2-utTrimer-ScDbp1p(2-154)-mTagBFP2-PDZ-tPGK1 | yKS101 | pKS1333 | HIS3 | 1d | This Study |
| yKS403 | yKS101 X-4:: pTEF1-Venus-ScDbp1p(2-154)mut-ScFis1p(119-155)!-tSSA1-Con1-pHHF2-utTrimer-ScDbp1p(2-154)-mTagBFP2-PDZ-tPGK1 | yKS101 | pKS1334 | HIS3 | 1d | This Study |
| yKS380 | CEN.PK2-1C X-4:: pTEF1-ALFA_Nb-Crk_SH3-PTB-Venus-ScDbp1p(2-154)-ScUbc6p(233-250)!-tSSA1-Con1-pHHF2-utTrimer-ScDbp1p(2-154)-mTagBFP2-PDZ-tPGK1 | CEN.PK2-1C | pKS1126 | N/A | 2b, ED3a | This Study |
| yKS381 | CEN.PK2-1C X-4:: pTEF1-ALFA_Nb-Crk_SH3-PTB-Venus-ScDbp1p(2-154)-ScGap1p(550-602)!-tSSA1-Con1-pHHF2-utTrimer-ScDbp1p(2-154)-mTagBFP2-PDZ-tPGK1 | CEN.PK2-1C | pKS1262 | N/A | 2b, ED3a | This Study |
| yMW941 | CEN.PK2-1C OYE2:: loxP-pGPD-NsrR-ENO2t-loxP; OYE3:: loxP-pGPD-HygR-TDH1t-loxP | CEN.PK2-1C |  | HygR, NatR |  | Ref <sup>25</sup> |
| yKS109 | yMW941 XI-1:: pTDH3-EfMvaE-tENO1-Con1-pTEF1-Scldi1p-tADH1-Con2-pPGK1-ScErg20p(F96W_N127W)-tSSA1-Con3-pCCW12-ObGES-tPGK1 | yMW941 | pKS557 | HygR, NatR |  | This Study |
| yKS404 | yKS109 XII-5:: pGAL1-CrG8H-PDZL-tENO1-Con1-Spacer-Con2-Spacer-Con3-pGAL10-CrCPR-PTBL-tPGK1-Con4-pHTB2-SH3L-CrCYB5-tTDH1 | yKS109 | pKS1337 | HygR, NatR | 3b | This Study |
| yKS228 | yKS109 XII-5:: pGAL1-CrG8H-SH3L-tENO1-Con1-Spacer-Con2-Spacer-Con3-pGAL10-CrCPR-PTBL-tPGK1-Con4-pHTB2-ALFAL-CrCYB5-tTDH1 | yKS109 | pKS1003 | HygR, NatR | 3b, 3g | This Study |
| yKS231 | yKS109 XII-5:: pGAL1-CrG8H-dSH3L-tENO1-Con1-Spacer-Con2-Spacer-Con3-pGAL10-CrCPR-dPTBL-tPGK1-Con4-pHTB2-ALFAL-CrCYB5-tTDH1 | yKS109 | pKS1077 | HygR, NatR | 3b | This Study |
| yKS122 | CEN.PK2-1C XI-1:: pTDH3-EfMvaE-tENO1-Con1-pTEF1-Scldi1p-tADH1-Con2-pPGK1-ScErg20p-tSSA1-Con3-pCCW12-GB1-AaADS-tPGK1 | CEN.PK2-1C | pKS709 | N/A |  | This Study |
| yKS342 | yKS122 LEU2:: pTEF2-AaADH1-tENO2-Con1-pHHF2-AaALDH1-tTDH1 | yKS122 | pKS1304 | LEU2 |  | This Study |
| yKS406 | yKS342 XII-5:: pGAL1-AaCYP71AV1-PDZL-tENO1-Con1-pGAL10-AaCPR-PTBL-tPGK1-Con2-pHTB2-WH1L-AaCYB5-tTDH1 | yKS342 | pKS1324 | LEU2 | 3b | This Study |
| yKS407 | yKS342 XII-5:: pGAL1-AaCYP71AV1-SH3L-tENO1-Con1-pGAL10-AaCPR-PTBL-tPGK1-Con2-pHTB2-WH1L-AaCYB5-tTDH1 | yKS342 | pKS1266 | LEU2 | 3b | This Study |
| yKS409 | yKS342 XII-5:: pGAL1-AaCYP71AV1-dSH3L-tENO1-Con1-pGAL10-AaCPR-dPTBL-tPGK1-Con2-pHTB2-WH1L-AaCYB5-tTDH1 | yKS342 | pKS1269 | LEU2 | 3b | This Study |
| EBY.VW4000 | CEN.PK2-1C hxt13Δ::loxP hxt15Δ::loxP hxt16Δ::loxP hxt14Δ::loxP hxt12Δ::loxP hxt9Δ::loxP hxt11Δ::loxP hxt10Δ::loxP hxt8Δ::loxP hxt514Δ::loxP hxt2Δ::loxP stl1Δ::loxP agt1Δ::loxP ydl247wΔ::loxP yjr160cΔ::loxP | CEN.PK2-1C |  | N/A<br>(Unable to use glucose) |  | Ref <sup>67</sup> |
| yKS134 | EBY.VW4000 XI-1:: pTDH3-ScTkl1p-tENO1-Con1-pCCW12-ScTal1p-tSSA1-Con2-pPGK1-ScRpi1p-tADH1-Con3-pHHF2-ScRpe1p-tPGK1 | EBY.VW4000 | pKS803 | N/A |  | This Study |
| yKS221 | yKS134 X-2:: pCCW12-ALFAL-ScHxt7p-tENO1 | yKS134 | pKS1043 | N/A |  | This Study |
| yKS244 | yKS221 XII-5:: Spacer-Con1-pTEF1-SH3L-PsXyl1!-tADH1-Con2-pHHF2-PTBL-PsXyl2!-tSSA1-Con3-pTDH3-WH1L-PsXyl3!-tENO1 | yKS221 | pKS1068 | N/A | 4b, 4d | This Study |
| yKS245 | yKS221 XII-5:: Spacer-Con1-pTEF1-SH3L-PsXyl1(R276H)!-tADH1-Con2-pHHF2-PTBL-PsXyl2!-tSSA1-Con3-pTDH3-WH1L-PsXyl3!-tENO1 | yKS221 | pKS1070 | N/A | 4b, 4d | This Study |
| yKS329 | yKS221 XII-5:: Spacer-Con1-pTEF1-dSH3L-PsXyl1(R276H)!-tADH1-Con2-pHHF2-dPTBL-PsXyl2!-tSSA1-Con3-pTDH3-WH1L-PsXyl3!-tENO1 | yKS221 | pKS1075 | N/A | 4b, 4d | This Study |
| yKS331 | CEN.PK2-1C 2μ: pGAL1-utTrimer-ScDbp1p(2-154)-mCherry-PTB-2xCrk_SH3-tENO1 | CEN.PK2-1C | pKS1241 | LEU2 | ED6b | This Study |

**Supplementary Video 1. Slab Dbp1n condensates partition NADH.** In residue-resolution coarse-grained molecular dynamics simulation of biomolecular condensates composed of Dbp1n (yellow) with slab geometry, NADH (green) strongly partitions into the protein dense phase.

**Supplementary Video 2. In slab FUSn condensates, the NADH partition coefficient is less than in Dbp1n condensates.** Using residue-resolution coarse-grained molecular dynamics simulation of biomolecular condensates composed of FUSn (yellow) with slab geometry, NADH (green) is not strongly partitioned into the protein dense phase.

**Supplementary Video 3. Dbp1n sMACs partition NADPH.** In residue-resolution coarse-grained molecular dynamics simulation of sMACs composed of Dbp1n (green), anchored to a membrane-bound organelle (gray), where the condensate strongly partitions NADPH (red).

**Supplementary Video 4. In FUSn sMACs, the NADPH partition coefficient is less than in Dbp1n condensates.** Residue-resolution coarse-grained molecular dynamics simulation of sMACs composed of FUSn (green), anchored to a membrane-bound organelle (gray). The condensate partitions NADPH (red) more weakly than the corresponding simulation where the FUSn IDR is replaced with Dbp1n.

